# A Diverse Range of Factors Affect the Nature of Neural Representations Underlying Short-Term Memory

**DOI:** 10.1101/244707

**Authors:** A. Emin Orhan, Wei Ji Ma

## Abstract

Sequential and persistent activity models are two prominent models of short-term memory in neural circuits. In persistent activity models, memories are represented in persistent or nearly persistent activity patterns across a population of neurons, whereas in sequential models, memories are represented dynamically by a sequential pattern of activity across the population. Experimental evidence for both types of model in the brain has been reported previously. However, it has been unclear under what conditions these two qualitatively different types of solutions emerge in neural circuits. Here, we address this question by training recurrent neural networks on several short-term memory tasks under a wide range of circuit and task manipulations. We show that sequential and nearly persistent solutions are both part of a spectrum that emerges naturally in trained networks under different conditions. Fixed delay durations, tasks with higher temporal complexity, strong network coupling, motion-related dynamic inputs and prior training in a different task favor more sequential solutions, whereas variable delay durations, tasks with low temporal complexity, weak network coupling and symmetric Hebbian short-term synaptic plasticity favor more persistent solutions. Our results help clarify some seemingly contradictory experimental results on the existence of sequential vs. persistent activity based memory mechanisms in the brain.

## Introduction

Short-term memory is a fundamental cognitive function for both humans and other animals. Despite its importance, its neural basis largely remains an open problem. The classical view of how a short-term memory might be implemented in the brain relies on the idea of a fixed point attractor [1, 2]. In this view, a memory is maintained via persistent activity of individual neurons. By virtue of their persistent activity, those neurons continue to represent information in the absence of any sensory stimulation. However, persistent activity of individual neurons is not necessary for maintaining information in short-term memory; dynamic activity patterns can also maintain short-term memories [3-5]. According to this alternative view, individual neurons can be active only transiently, while the population as a whole maintains the memory through a dynamically changing activity pattern across time.

It has been an ongoing debate whether one of these alternative pictures provides a more accurate representation of the neural mechanism (or mechanisms) underlying short-term memory than the other one [6, 7]. Experimental evidence for both alternatives has been reported previously: for example, [8-12] observed persistent or nearly persistent activity during the delay period of short-term memory tasks, whereas [13-18] observed sequential or dynamic activity patterns. These studies used different tasks, different stimuli, different experimental designs and sometimes recorded from different areas or even from different species. It is difficult to know which of these differences might be relevant for the observed differences in the mnemonic activity patterns. Although this question can, in principle, be addressed experimentally by running many experiments, systematically varying each experimental factor or neural circuit property that could conceivably have an effect on the observed differences, this would be too costly. Instead, here we address this question by performing these experiments *in silico*. This allows us to not only identify the relevant factors, but also understand mechanistically why those factors have the effects that they do.

More specifically, we trained recurrent neural networks on a range of short-term memory tasks and investigated the effects of a diverse array of task-related and circuit-related factors on the sequentiality or persistence of the emergent activity patterns: (i) the task; (ii) other experimental variables such as delay duration variability or whether the task had a navigation component; (iii) whether the network was previously trained on another task; (iv) intrinsic network properties such as the intrinsic timescale of individual neurons and the strength of coupling between the neurons; and (v) Hebbian short-term synaptic plasticity.

We find that sequential and nearly persistent solutions are both part of a spectrum that emerges naturally in trained networks under different conditions. Tasks with higher temporal complexity, fixed delay durations, stronger network coupling between the neurons, prior training in another task and task-irrelevant motion-related dynamic cues that arise in navigation-like tasks all increase the sequentiality of the emergent solutions; whereas tasks with lower temporal complexity, variable delay durations, weak coupling between the neurons and symmetric Hebbian short-term synaptic plasticity reduce the sequentiality of the solutions. Furthermore, having complete access to the networks and their behavior allowed us to develop a detailed mechanistic understanding of the circuit mechanism that generates sequential vs. persistent mnemonic activity and why the aforementioned factors have the effects that they do on the sequentiality or persistence of the neural responses.

## Results

### Experimental setup

#### Networks

In our main simulations, we used vanilla recurrent neural networks with rectified linear recurrent units (Figure 1a; see *Methods*). The input to the network was provided in the form of a population of Poisson neurons, emitting independent Poisson counts at each time step of the simulation. Experimental evidence suggests that both intrinsic time constants of individual neurons and the overall coupling strength between them can vary significantly across cortex [19, 20]. In order to tease apart the potential effects of these two factors, we initialized the recurrent connectivity matrix as *λ*_0_**I** + *σ*_0_Σ_off_ where *λ*_0_ and *σ*_0_ are hyperparameters controlling the amount of initial self-recurrence and recurrence from the rest of the network respectively, **I** is the identity matrix and Σ_off_ is an off-diagonal matrix whose off-diagonal entries are drawn independently from a zero-mean normal distribution with standard deviation 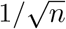 (Figure 2a), where *n* is the number of recurrent units in the network. Since regularization of the network parameters (or the recurrent activity) can sometimes significantly impact the nature of the emergent solutions [21-23], we also placed an *l*_2_-norm regularizer on the network parameters throughout training and controlled its strength through another hyperparameter, *ρ*. We repeated our main simulations for 800 different hyperparameter configurations drawn over a grid in the (*λ*_0_*, σ*_0_*, ρ*) space. On this grid, *λ*_0_ took 10 uniformly-spaced values between 0.8 and 0.98, *σ*_0_ took 10 uniformly-spaced values between 0 and 0.4025 and *ρ* took 7 logarithmically-spaced values between 10^−6^ and 10^−3^ as well as *ρ* = 0. In general, we chose these ranges to be as large as possible subject to the trainability of the networks, such that values outside of these ranges, in general, significantly impeded the trainability of the networks. These choices still gave rise to a wide range of initial network dynamics, from quickly decaying to strongly unstable (Supplementary Figure S1). Qualitatively, increasing *λ*_0_ has the effect of increasing the intrinsic time constant of the individual neurons, making their activity more persistent in response to an input pulse. Increasing *σ*_0_, on the other hand, introduces oscillatory components to the network response.

**Figure 1:**
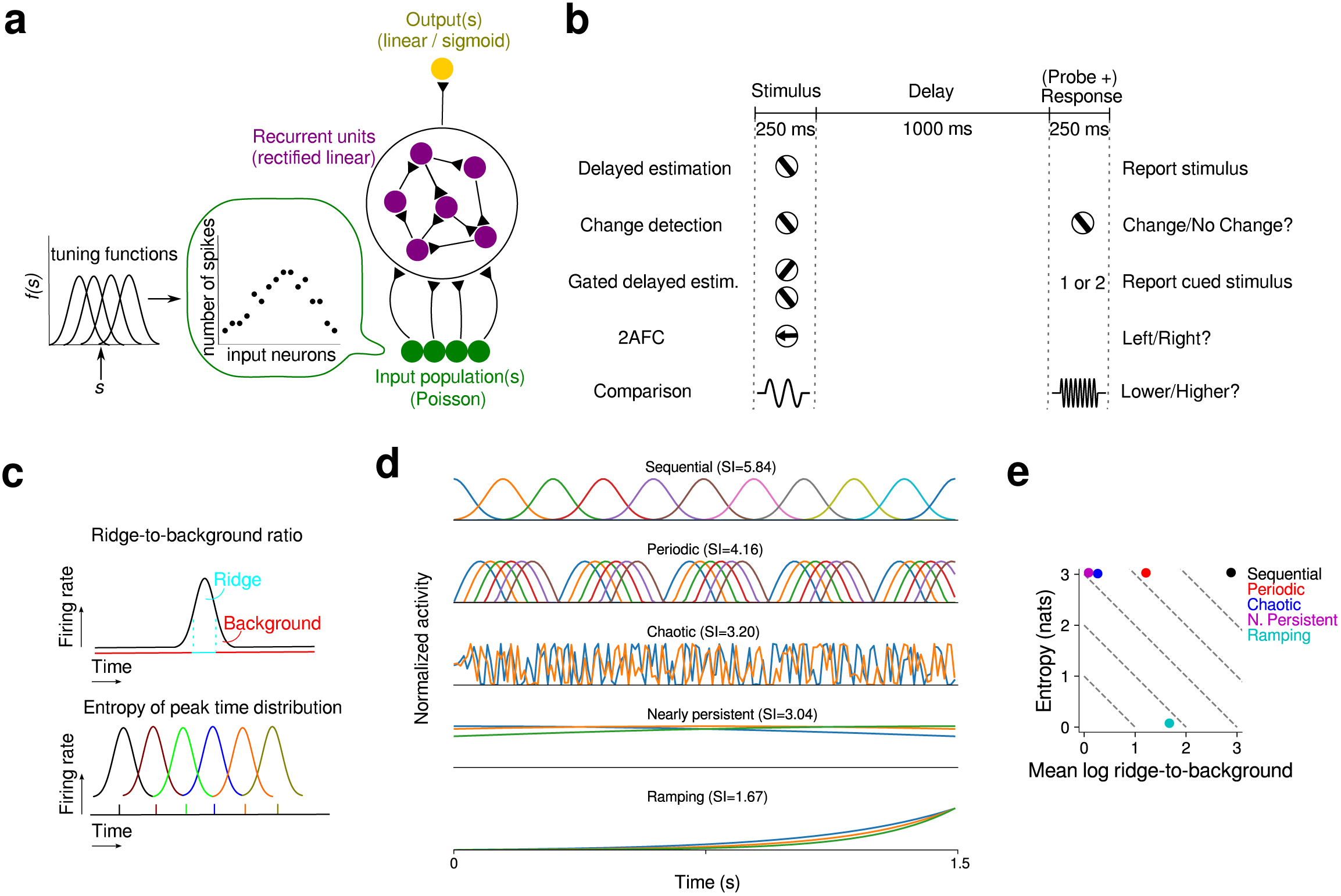
Experimental setup. a Schematic diagram of recurrent networks. The input neurons are Poisson neurons providing noisy information about the stimulus or the stimuli. These neurons project onto the recurrent neurons which are modeled as rectified linear units (ReLUs). Recurrent neurons, in turn, project onto the output unit or units, which are either linear or sigmoidal in different tasks. b The five main experimental tasks and the common trial structure. c Two factors determining the sequentiality index (SI): the ridge-to-background ratio [16] measures the temporal localization of the activity of individual units; the entropy of the peak time distribution measures the uniformity of the peak response times of the units in a given trial. The SI for a given trial is then given by the sum of the mean log ridge-to-background ratio of the recurrent units and the entropy of the peak time distribution. d Example idealized single-trial activity patterns with the corresponding sequentiality indices (SI) indicated at the top of each panel. Different colors represent the temporal activity patterns of a subset of individual units. These example trials were generated with the same number of recurrent units and time steps as in the simulations in the rest of the paper. Hence, the SI values here are directly comparable to the SI values reported elsewhere in the paper. A small amount of noise, independent across neurons and time, was added to the responses of all neurons in order to break possible ties in determining peak response times. **e** shows how the example trials shown in d score along each of the two dimensions defining the SI. Dashed lines represent several iso-SI contours. All examples except for the ramping one score close to maximum on the entropy dimension, hence their SIs are largely distinguished by the mean ridge-to-background ratio. Note that the nearly persistent example was generated by broadening the temporal activity profiles in the sequential example. It has thus the same peak time entropy as the sequential example, but has a much smaller mean ridge-to-background ratio. The ramping example, on the other hand, has minimal peak time entropy and a medium mean ridge-to-background ratio.

#### Tasks

In order to eliminate potential differences due to trial structure, we used a common trial structure for all our tasks (Figure 1b). Each trial started with the presentation of one or two stimuli for 250 ms. A delay period of 1000 ms then followed. After the delay, there was a response period of 250 ms. In some tasks, a second stimulus or a cue appeared during the response period, in which case the target response depended on this second stimulus or cue.

We considered five main tasks in our experiments (Figure 1b; see *Methods* for task details): (i) delayed estimation with one stimulus (DE-1) or with two stimuli (DE-2), where the task was to report the stimulus or stimuli presented at the beginning of the trial; (ii) change detection (CD), where the task was to report whether the stimulus presented before the delay was the same as the stimulus presented after the delay (e.g. [24]); (iii) gated delayed estimation (GDE), where two stimuli were presented simultaneously at the beginning of the trial and the task was to report the cued one after the delay (e.g. [24]); (iv) 2AFC, where one of two possible stimuli (e.g. left vs. right moving dots) was presented at the beginning of the trial and the task was to report which one was presented (e.g. [11, 16]); (v) comparison (COMP), where the task was to report whether the stimulus presented before the delay was smaller or larger than the one presented after the delay (e.g. [10]).

#### Quantifying sequentiality

Intuitively, there are two requirements for the recurrent activity of a population of neurons to be considered sequential (Figure 1c): (i) each neuron should be active only during a short interval compared to the duration of the trial; (ii) the active periods of the neurons should tile the entire duration of the trial approximately uniformly. Thus, we designed a *sequentiality index* (SI) that takes into account both of these desiderata. The sequentiality index for a given trial is defined as the sum of the entropy of the peak response time distribution of the recurrent neurons and the mean log ridge-to-background ratio of the neurons, where the ridge-to-background ratio for a given neuron is defined as the mean activity of the neuron inside a small window around its peak response time divided by its mean activity outside this window [16]. The sequentiality index for a given experimental condition is then determined by averaging over the sequentiality indices of all trials belonging to that condition. Figure 1d shows some idealized single-trial temporal activity patterns and the corresponding SIs. These examples were generated using the same number of recurrent neurons and time steps as in other simulations in this paper, hence the SI values reported in Figure 1d are directly comparable to those reported elsewhere in the paper.

### Factors affecting the sequentiality of the responses

#### Intrinsic circuit properties affect sequentiality

In successfully trained networks, the sequentiality of the recurrent activity increased with the initial network coupling, *σ*_0_ (Figure 2b); it did not change significantly with the initial intrinsic timescale of individual units, *λ*_0_ (Figure 2c) and it slightly but significantly decreased with the regularization coefficient *ρ* (Figure 2d). Larger *σ*_0_ values introduce higher frequency oscillatory dynamics in the initial network, which promotes the emergence of high frequency sequential structure in the trained networks. Larger *ρ* values, on the other hand, have the opposite effect.

**Figure 2:**
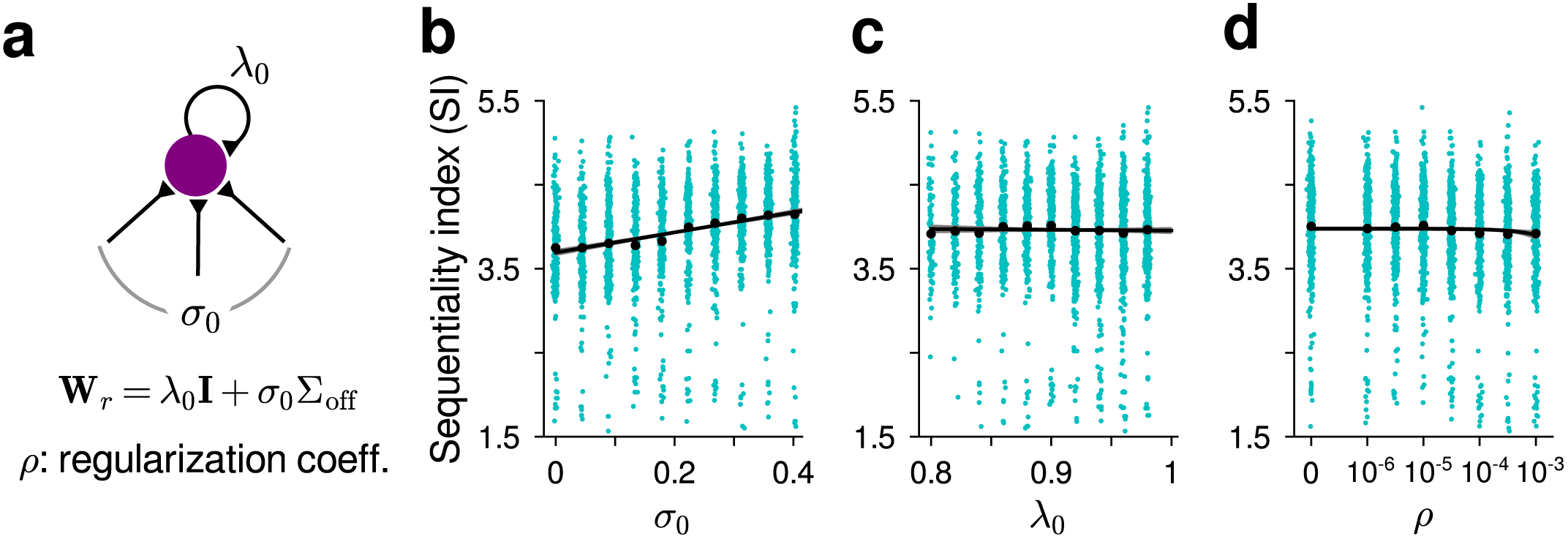
Intrinsic circuit properties and their effect on the sequentiality of the recurrent activity in trained networks. a The recurrent connectivity matrix was initialized as **W**_*r*_ = *λ*_0_**I** + *σ*_0_Σ_off_ where *λ*_0_ controls the initial intrinsic timescale of individual units and *σ*_0_ controls the size of the initial coupling between the units. We also varied the strength of the *l*_2_-norm regularization on the parameters, controlled by the coefficient *ρ*. Our basic experiments were repeated with 800 different *λ*_0_, *σ*_0_, *ρ* values on a 10 × 10 × 8 grid over the three-dimensional hyperparameter space (*λ*_0_, *σ*_0_, *ρ*). b SI increased with *σ*_0_ (linear regression slope: 1.20, *R*^2^ = 0.08, *p* < 10^−53^). c SI did not change significantly with *λ*_0_ (*p* = 0.64). d SI slightly decreased with *ρ* (linear regression slope: −76, *R*^2^ = 0.002, *p* < .05). Each cyan dot corresponds to the mean SI for a particular hyperparameter setting and a particular task. Black dots represent the means. Solid black lines are the linear fits and shaded regions are 95% confidence intervals for the linear regression (confidence intervals are usually too small to be clearly noticeable on the plotted scale).

#### Temporal complexity of tasks affects sequentiality

There was significant variability in SI among the tasks (Figure 3a; see Supplementary Figures S2-S6 for example trials from all tasks under different hyperparameter settings). Indeed, task was the most predictive variable in a linear regression analysis of the SI including the variables task (coded ordinally) and the three hyperparameters *σ*_0_, *λ*_0_ and *ρ*: task alone yielded *R*^2^ = 0.20 compared to *R*^2^ = 0.08 for the next most predictive variable, *σ*_0_. Some tasks such as comparison or change detection led to highly sequential responses, whereas other tasks such as the basic 2AFC task led to less sequential and more persistent responses (Figure 3b). We hypothesized that this variability was related to the temporal complexity of the target functions that need to be learned in different tasks, where the target function complexity can be formalized as the mean temporal frequency of the target function [25], for instance. In change detection, gated delayed estimation and comparison tasks, the target function depends on the probe (or cue) stimulus presented after the delay period. Thus, these tasks have higher temporal complexity. In delayed estimation and 2AFC tasks, on the other hand, no probe is presented after the delay and the target response does not depend on what happens after the delay. Therefore, these tasks have lower temporal complexity. Implementing temporally more complex target functions requires higher frequency temporal basis functions and sequential activity in the recurrent population provides such a high frequency temporal basis.

**Figure 3:**
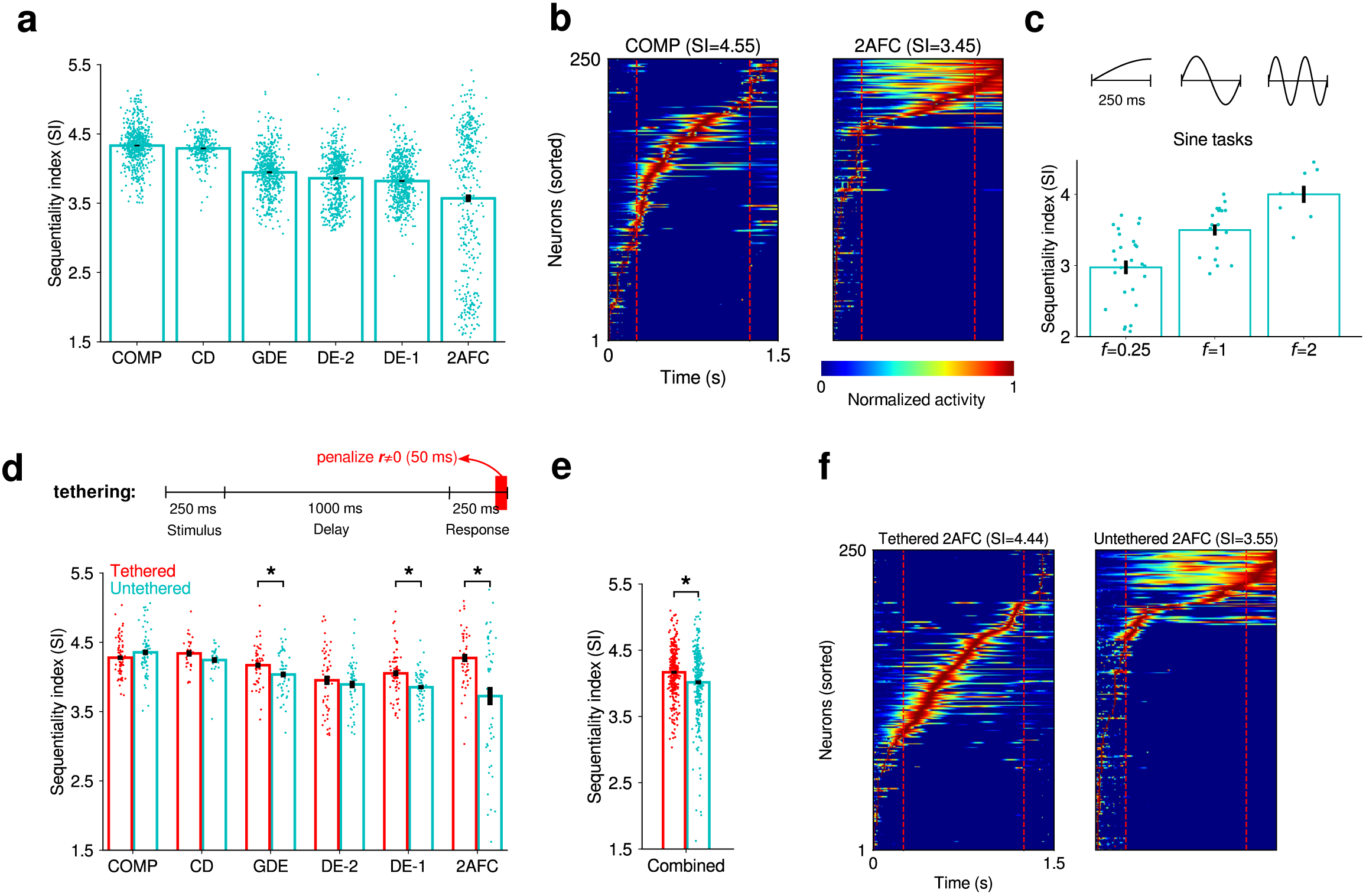
Temporal complexity of the task increases the sequentiality of the recurrent activity in trained networks. **a** Sequentiality index (SI) in different tasks. DE-1 and DE-2 refer to delayed estimation tasks with one stimulus and with two stimuli, respectively. Each dot corresponds to the mean SI for a particular setting of the hyperparameters. Error bars represent standard errors across different hyperparameter settings. **b** Normalized responses of recurrent units in a pair of example trials from the COMP and 2AFC tasks respectively, trained under the same hyperparameter setting. The SI values of the trials are indicated at the top of the corresponding panels. We chose representative trials with SI values close to the mean SI for the two tasks. Only the responses of the most active 250 units are shown here. The actual networks always consisted of 500 recurrent units. The remaining units were mostly or completely silent throughout the trial. **c** SI in the sine tasks. In these tasks, the network was trained to output a sine function with temporal frequency *f* during the response period (target functions are shown in the upper panel). Higher frequency target functions (larger *f*) led to more sequential responses: linear regression of SI on *f* yielded a slope of 0.60 (*R*^2^ = 0.43, *p* < 10^−7^). **d** Tethering manipulation and its effect on the SI of different tasks. Asterisk (*) indicates a significant difference at the *p* < .05 level (Welch’s *t*-test). **e** SI in the tethered vs. untethered conditions, combined across all tasks in **d**. **f** Normalized responses of recurrent units in a pair of example trials from the tethered and untethered versions of the 2AFC task respectively, trained under the same hyperparameter setting. We again chose representative trials with SI values close to the mean SIs of the two conditions.

To test this hypothesis more directly, we conducted two simple experiments. First, we trained networks to output sine functions with different temporal frequencies during the response period (upper panel in Figure 3c). The target function thus had the following form: sin(2*πf t*/*T*_resp_), where 0 ≤ *t* ≤ *T*_resp_ and *T*_resp_ denotes the duration of the response period. The networks received one-dimensional random input throughout the trial in these tasks. According to our hypothesis, target functions with higher temporal frequency (larger *f*) should lead to more sequential responses. We observed that this was indeed the case (Figure 3c): linear regression of SI on *f* yielded a slope of 0.60 (*R*^2^ = 0.43, *p* < 10^−7^).

Secondly, we introduced a “tethering” manipulation in our experimental design that increased the temporal complexity of the tasks. In tethered conditions, we put a strong penalty on recurrent responses deviating from 0 during the last 50 ms of the trial (upper panel in Figure 3d). An analogous tethering manipulation can be induced experimentally, for example, by optogenetic silencing of a relevant neural circuit toward the end of the trial. Tethering increases the temporal complexity of the task, because it forces the network’s output to sharply change from the roughly constant value it takes before the onset of tethering. We thus expected this manipulation to increase the sequentiality of the responses in successfully trained networks. Tethering indeed led to an overall increase in the sequentiality of the responses (Figure 3d-e). Importantly, in many cases, tethering changed the dynamics throughout the entire trial duration and not just toward the end of the trial (e.g. see the representative pair of trials in Figure 3f).

#### Hebbian short-term synaptic plasticity affects sequentiality

Short-term synaptic plasticity is a ubiquitous feature of synapses in real neural circuits [26]. A number of theoretical and experimental studies have suggested that short-term synaptic plasticity might be involved in short-term memory by storing information in an “activity-silent” format in synapses [27-29]. To investigate the effect of short-term synaptic plasticity on the sequentiality of the recurrent activity in our networks, we added a simple symmetric Hebbian short-term synaptic plasticity term to the recurrent weights (see *Methods*). This Hebbian contribution to the recurrent weights is sometimes known as “fast weights” in the machine learning literature [30].

Symmetric Hebbian short-term synaptic plasticity decreased the sequentiality of the recurrent activity in trained networks (Figure 4a). A symmetric contribution to the recurrent connectivity matrix reduces the high-frequency oscillatory dynamics in the network, which in turn reduces the sequentiality of the recurrent activity. We emphasize again the symmetry of the short-term synaptic plasticity rule considered here, since asymmetric associative rules (e.g. spike-timing-dependent plasticity) can often have opposite effects, as demonstrated in earlier studies [31-33]. We have tried several asymmetric variants of our Hebbian short-term synaptic plasticity rule, but we found these rules to be quite unstable in general and we were not able to train our networks successfully with these kinds of rules.

**Figure 4:**
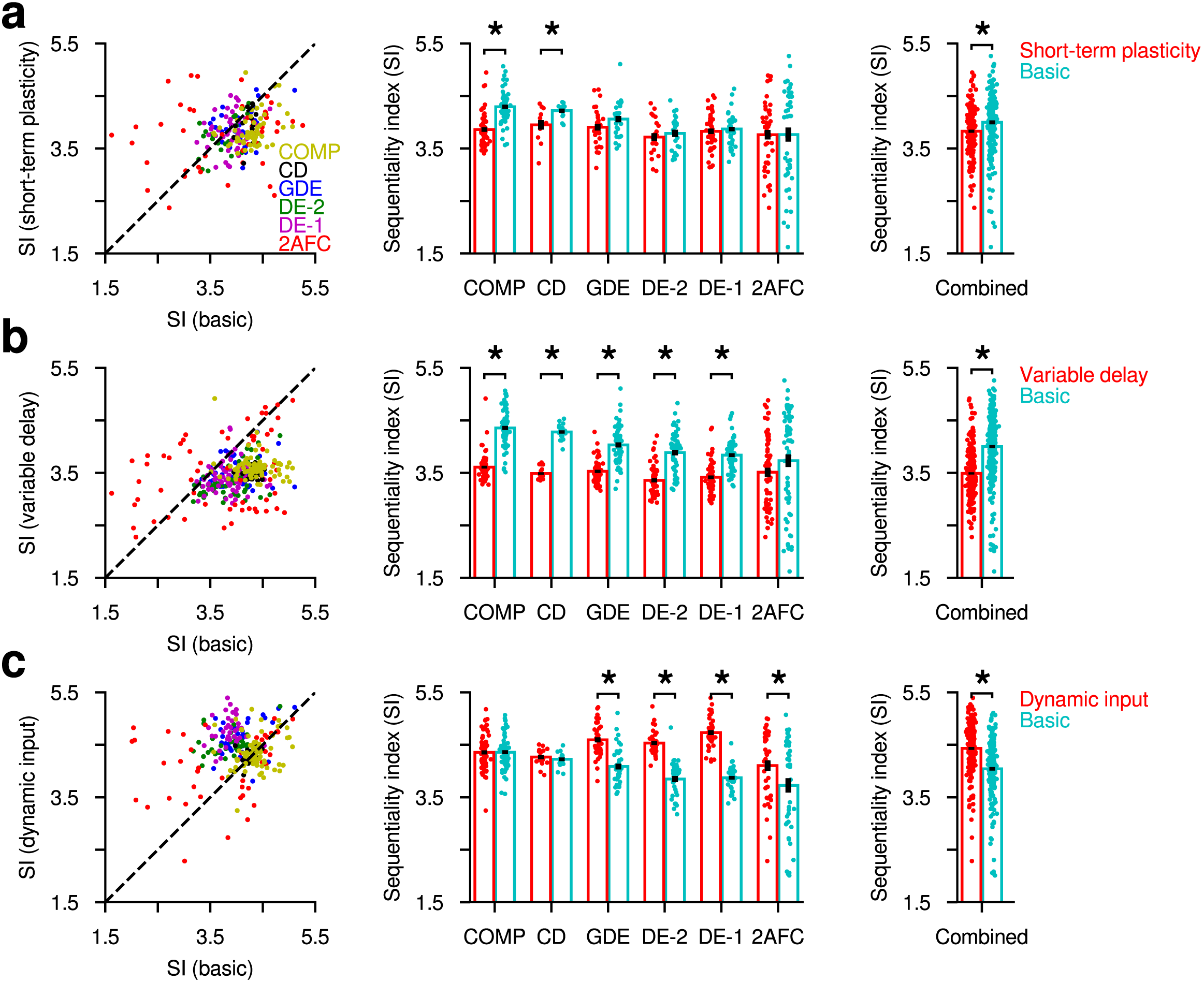
Hebbian short-term synaptic plasticity, delay duration variability and structured dynamic inputs affect the sequentiality of the recurrent activity in trained networks. **a** The effect of Hebbian short-term synaptic plasticity on the SI. The leftmost column shows a scatter-plot of the SI in the basic condition vs. the SI in the short-term plasticity condition. Each dot corresponds to a different initial condition and different colors represent different tasks. The middle column collapses the data across different initial conditions and compares the SI for each task. The rightmost column collapses the data further across tasks and compares the SI in the basic vs. short-term plasticity conditions for the combined data. Asterisk (*) indicates a significant difference at the *p* < .05 level (Welch’s *t*-test). Hebbian short-term synaptic plasticity decreased the SI. **b** The effect of delay duration variability on the SI. Delay duration variability decreased the SI. **c** The effect of structured dynamic input on the SI. Structured dynamic input increased the SI.

#### Delay duration variability affects sequentiality

Our simulations so far assumed a fixed delay duration of 1000 ms. However, experimenters sometimes use variable delay durations in short-term memory experiments. To test the effect of delay duration variability, we designed versions of each of our tasks with delay duration variability. In these versions, the delay duration was one of 100 ms, 400 ms, 700 ms, 1000 ms, chosen randomly on each trial. Variability in delay duration significantly decreased the sequentiality of the recurrent activity in successfully trained networks (Figure 4b). In sequential solutions, the representations of task-relevant variables change over time. Therefore, these representations cannot be decoded with a fixed decoder across time. However, the variable delay duration experiments demand that the learned representations be decodable with a fixed decoder at different delay durations, hence force the network to learn more stable representations across time.

#### Task-irrelevant structured dynamic inputs affect sequentiality

Motion-related signals animals receive during navigation-type experiments have previously been argued to be crucial for the generation of sequential neural activity observed in rodent experiments [34]. Our results from experiments without such motion-related signals clearly demonstrate that such signals are not necessary for the generation of sequential activity. However, it is still possible that because such signals already have a sequential structure, they may facilitate the generation of sequential activity in the network. To test this hypothesis, we designed navigation versions of our main experiments where, in each trial, the network was assumed to navigate through a linear track at constant speed. The network received noisy population coded information about its hypothetical location in the linear track, in addition to the task-relevant inputs it received (see *Methods*). The location information was irrelevant for performing the tasks, hence the network could safely ignore this information. These motion-related, task-irrelevant location signals significantly increased the sequentiality of the recurrent activity in successfully trained networks (Figure 4c), suggesting that the networks did not completely suppress these signals despite the fact that they were irrelevant to the tasks the networks were trained on.

#### Learning multiple tasks in sequence affects sequentiality

Our simulations so far assumed that each network is trained on a single task. However, a common situation that arises in many animal experiments is that the same animal may be trained on multiple tasks, usually sequentially. This can happen, for example, when an animal takes part in several different experiments throughout its lifetime, or when it learns to perform different tasks as part of a curriculum strategy for learning a more complex task. To investigate the effects of such sequential multi-task learning, we considered networks that learned a pair of tasks sequentially. We only considered the 2AFC–COMP and 2AFC–CD task pairs, trained in either order, because (i) these task pairs have the same number and type of inputs and outputs, hence do not require any changes in the network architecture and (ii) they have maximally different SIs when trained in isolation: the COMP and CD tasks have the largest SIs and the 2AFC task has the smallest SI among all tasks (Figure 3a). We then compared the SI in the second task of the pair with the SI of the same task when it was trained in isolation. Sequential multi-task training led to an overall increase in the SIs compared to the corresponding single task training conditions (Figure 5a-b). This might be expected in cases where the network was first trained on a high SI task and then on a low SI task (i.e. COMP→2AFC and CD→2AFC, although the effect was not significant in the latter case). More surprisingly, however, a significant increase in SI was also observed in the other direction, i.e. training in 2AFC→COMP produced a higher SI than training in COMP alone; and similarly, training in 2AFC→CD led to a higher SI than training in CD alone. We observed that this was because training a network in any task, including in low SI tasks such as 2AFC, consistently decreased the mean self-recurrence of the units, *λ* ≡ *(W_ii_)*, and increased the size of the fluctuations in the strength of recurrent coupling to the rest of the network, *σ* ≡ std(*W_ij,i/_*_=*j*_), compared to the initial weights (Figure 5c). Hence, for the second task in the pair, the effect of prior training in another task is analogous to an increase in the hyperparameter, *σ*_0_, which was shown to increase the SI above (Figure 2b).

**Figure 5:**
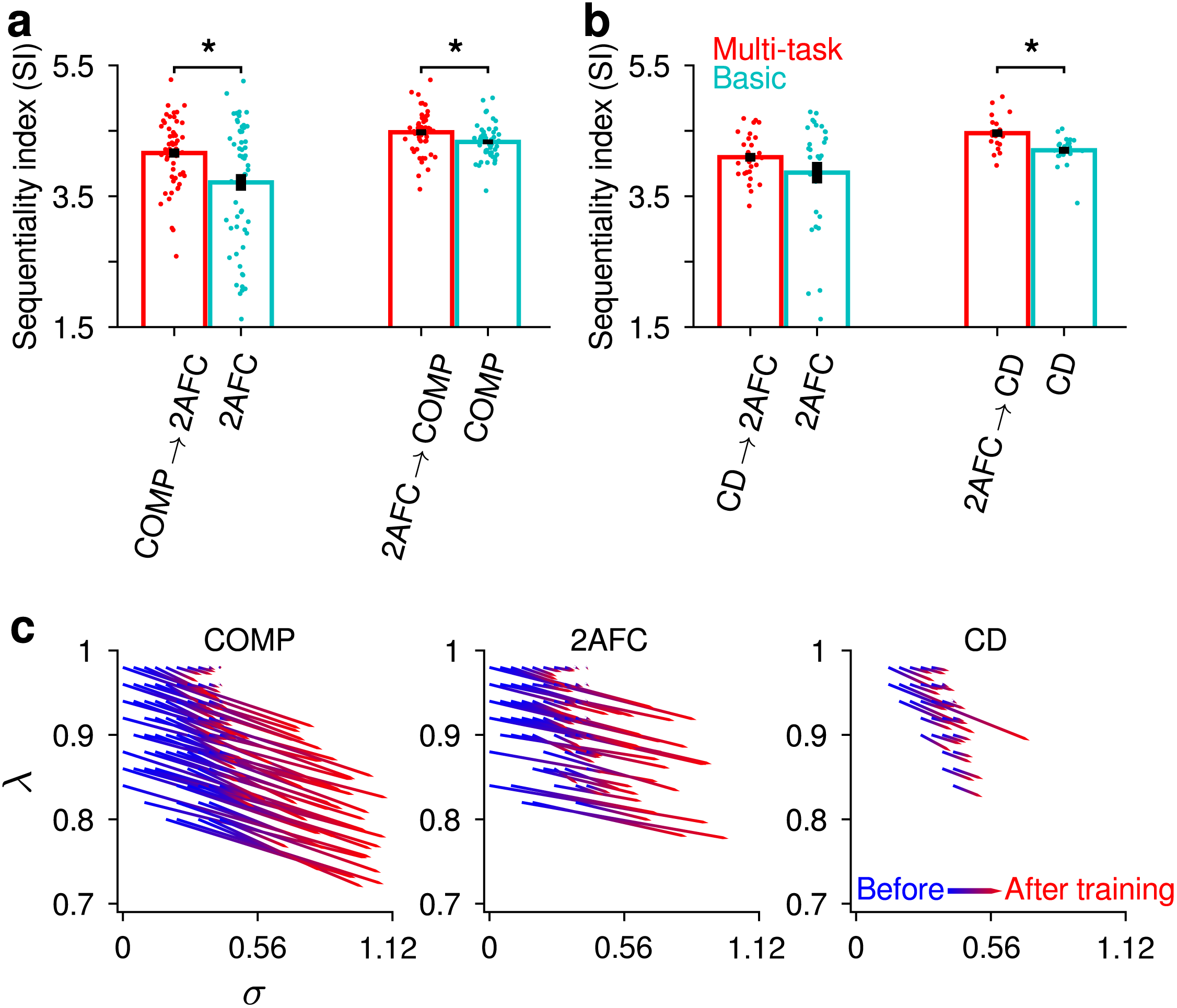
Multi-task learning experiments. **a** Results for the 2AFC–COMP task pair. **b** Results for the 2AFC–CD task pair. The red bars show the results for the multi-task training conditions, the cyan bars show the results for the corresponding single task training conditions. The right arrow indicates the order of training: e.g. “COMP 2AFC” means the network was first trained on the COMP task and then on the 2AFC task. Error bars represent standard errors across different hyperparameter settings. Asterisk (*) indicates a significant difference at the *p* < .05 level (Welch’s *t*-test). **c** Training in a task consistently reduces the mean self-recurrence, *λ* ≡ *(W_ii_)*, and increases the fluctuations in the strength of recurrent coupling to the rest of the network, *σ* ≡ std(*W_ij,i≠j_*). Note that *λ* = *λ*_0_ and *σ* = *σ*_0_ (as defined in Figure 2a) before training.

#### Circuit mechanism that generates sequential vs. persistent activity

To probe the circuit mechanism generating sequential vs. persistent activity in trained networks, we performed an analysis proposed by ref. [35]. In this analysis, we first ordered the recurrent neurons in the network by their time of peak activity. We then measured the mean and standard deviation of the recurrent weights, *W_ij_*, as a function of the order difference between two neurons, *i* − *j*. In trained networks, connections from earlier peaking neurons to later peaking neurons had, on average, larger weights than connections from later peaking neurons to earlier peaking neurons. The mean connection weight was an approximately monotonically increasing function of *i* − *j* (Figure 6a-b). This particular asymmetric structure generated sequential activity in the network with increasingly prolonged responses in later peaking neurons in the sequence (Figure 6e). However, in trained networks with high sequentiality index (SI > 5), a prominent asymmetric peak appeared in the connection weight profile (inset in Figure 6a). This asymmetric peak corresponds to strengthened connections between temporally close neurons in the sequence at the expense of weakened connections between temporally distant neurons, with connections in the “forward” direction being strengthened more than those in the opposite direction. This, in turn, led to more strongly sequential responses in the network (Figure 6d) by reducing the temporal smearing of the responses that took place in networks with low sequentiality index (SI < 2.5), which did not display such a peak in their connection weight profile (Figure 6b). A simplified model that only incorporated the nonlinearity and idealized versions of the mean connection weight profiles shown in Figure 6a-b captured essential aspects of the difference between the two cases (Supplementary Figure S7).

Importantly, the preceding analysis suggests that both sequential and persistent activity patterns underlying short-term memory in different conditions emerge as two ends of a spectrum in trained networks, rather than being categorically different solutions.

**Figure 6:**
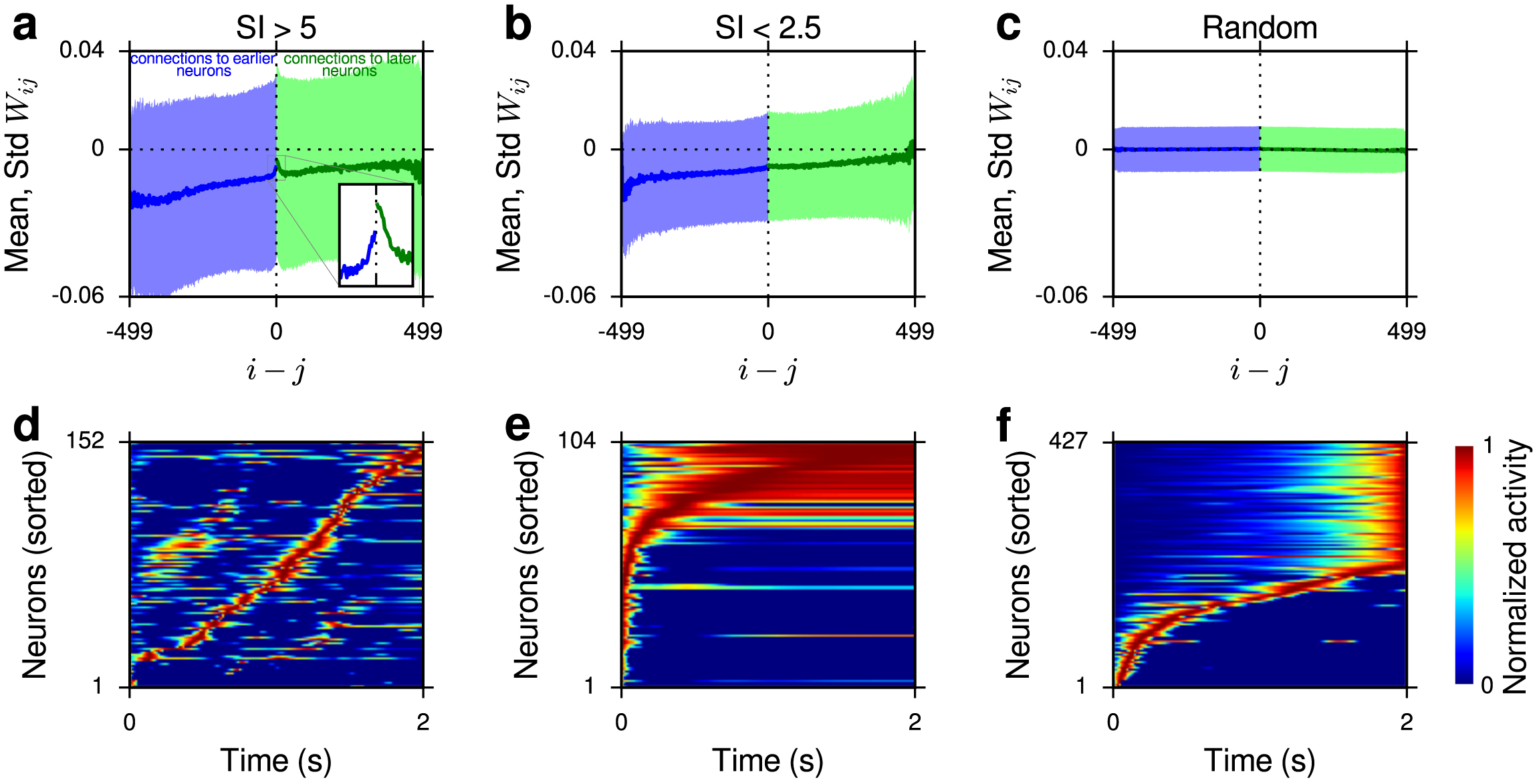
Circuit mechanism that generates sequential vs. persistent activity. **a, b, c** Neurons are first sorted by the time of their peak activity. We then plot the mean and standard deviation of the recurrent weights, *W_ij_*, as a function of the difference between the orders of the neurons in the sequence, *i j*. A positive *i j* value (green) indicates a connection from an earlier peaking neuron to a later peaking neuron. A negative *i j* value (blue) indicates a connection from a later peaking neuron to an earlier peaking neuron. Solid lines represent means and shaded regions represent standard deviations. **a** shows the results for all trained networks with SI > 5, **b** shows the results for all trained networks with SI < 2.5 and **c** shows the results for untrained random networks. The self-recurrence term corresponding to *i j* = 0 is not shown for clarity. **d, e, f** show normalized responses of neurons in example trials simulated with connectivity matrices drawn from the profiles shown in **a, b, c**, respectively (see *Methods* for details). Only the active neurons are shown in these plots.

### Robustness of the results to variations in some architectural and ex-perimental choices

In our simulations thus far, we have used recurrent networks of rectified linear units (ReLUs). This particular nonlinearity is unbounded on one side, hence it may be considered biologically unrealistic, even though in trained networks the recurrent units typically did not attain unrealistically large values. It is thus important to check whether our main results still hold for a nonlinearity saturating on both sides. For this purpose, we reproduced our main experiments with a simple modification to the networks, namely we replaced the ReLU nonlinearity with a clipped version of it that was bounded above by a maximum value (which we chose to be 100). Overall, the results from these simulations were qualitatively in agreement with the results from the ReLU networks. In particular, the hyperparameters *σ*_0_ and *ρ* (but not *λ*_0_) had similar effects on SI, the ordering of the tasks by SI was similar and the underlying mechanism that generated more sequential vs. more persistent activity in different conditions was also similar in the clipped ReLU networks (see Supplementary Figure S8).

Secondly, in our simulations, we chose the input noise levels to be roughly consistent with those used in ref. [36], where generic neural networks were trained on tasks similar to those considered here in psychophysically realistic input noise regimes. To investigate the sensitivity of our results to the amount of input noise, we re-ran our main experiments with up to 2.5 times lower and up to 2 times higher levels of input noise. Increasing the input noise slightly increased the SI (Supplementary Figure S9c). Importantly, even when we restricted the analysis to the lowest and the highest levels of input noise, we observed qualitatively very similar results to those reported above for our main experiments: i.e. the hyperparameters *σ*_0_ and *λ*_0_ had similar effects on SI, the ordering of the tasks by SI was similar and the circuit mechanism generating more sequential vs. more persistent solutions under different conditions was also similar (Supplementary Figures S10-S11).

## Discussion

We have identified a diverse range of circuit-related and task-related factors affecting the sequentiality or persistence of recurrent neural activity underlying short-term memory maintenance. Tasks with higher temporal complexity, fixed delay durations, stronger network coupling between neurons, motion-related dynamic cues and prior training in other tasks promote more sequential activity in trained networks; whereas tasks with lower temporal complexity, variable delay durations, weak coupling between neurons and symmetric short-term synaptic plasticity promote more persistent activity.

We have also developed a detailed mechanistic understanding of the circuit mechanism that generates sequential vs. persistent activity. In all trained networks, the basic mechanism implementing short-term memory maintenance is sequential recurrent activity generated by a non-normal recurrent connectivity matrix (see Supplementary Figure S12 for Schur decompositions of trained recurrent connectivity matrices), with increasingly prolonged responses as the activity travels along the sequence. In networks with more sequential activity, however, this temporal smearing is reduced by a characteristic asymmetric peak in the weight profile that corresponds to strengthened connections between temporally close neurons in the sequence (at the expense of weakened connections between temporally distant neurons), with connections in the “forward” direction being preferentially strengthened (Figure 6).

An important question to consider is why trained networks develop a short-term memory maintenance mechanism that relies on non-normal recurrent dynamics, even when the recurrent connectivity is initialized close to a normal matrix. For linear networks, it has been previously shown that any dynamical system with optimal memory properties must be non-normal and a feedforward chain is one of the simplest examples of such a non-normal dynamical system [41]. However, there are important differences between our networks and the simplified setup studied in [41]. Therefore, it remains to be seen whether this previous work can explain the emergence of non-normal structures in our trained networks. Another possibility is that non-normal solutions may just be more generic than normal solutions so that a randomly initialized network is more likely to converge to a non-normal solution.

A previous work (ref. [35]) also investigated the circuit mechanism underlying the generation of sequential activity in recurrent neural networks. However, that work did not train the networks to perform any short-term memory task, but rather trained them explicitly to generate sequential activity. Our work, on the other hand, shows that sequential activity emerges naturally in networks trained to perform short-term memory tasks and certain factors identified here facilitate the emergence of such sequential activity.

Rajan et al. [35] discovered qualitatively different mechanisms generating sequential activity as the fraction of trainable connections was varied in their networks. When only a small fraction of the connections were trainable, they found an input-dependent mechanism for the generation of sequences that is different from the mechanism uncovered in this work. Our mechanism relies on an asymmetric recurrent connectivity matrix and is conceptually similar to the sequence generation mechanism they found in networks where all connections were trainable. The particular asymmetry we found, however, is qualitatively different from the one found in their work. This difference is largely due to the difference in the training signals (our networks were trained on actual short-term memory tasks without constraining the dynamics, theirs were trained to generate sequential activity). Training the networks to explicitly generate sequential activity constrains the recurrent connectivity more strongly and results in more structured weight profiles, especially with the tanh nonlinearity used in [35] (Supplementary Figure S13).

In addition to the difference in training signals, there are two other differences between [35] and our work. First, they used tanh units, whereas we used ReLUs in our networks. We were not able to successfully train networks of tanh units in any of our tasks, neither with the particular initialization we used, nor with more standard initializations. However, we reproduced our experiments with two other activation functions, exponential linear [42] and softplus [43] (in addition to the double-sided saturating clipped ReLU nonlinearity discussed above), and found asymmetries in the trained recurrent connectivity matrices that were qualitatively similar to those observed in our ReLU networks (Supplementary Figure S14). Second, Rajan et al.’s networks always received dynamic inputs, whereas in our basic condition, the networks did not receive any input during the delay (except for a very small amount of spontaneous input due to the stochasticity of input units; see *Methods*). Hence their simulations were more similar to our dynamic, motion-related input condition than to our basic condition. Together, Rajan et al. [35] and this study demonstrate a multiplicity of ways in which sequential activity can be generated in neural circuits.

Our results concerning the various factors affecting the sequentiality or persistence of neural activity underlying short-term memory immediately lead to experimental predictions that can be tested in the lab. There is already experimental evidence confirming the effects of some of these factors. For instance, [11] observed more persistent responses in mouse posterior parietal cortex than [16] did in the same area while animals in both studies were performing visual short-term memory tasks. There were, however, crucial differences between the experimental designs in these studies: in [11], the task was not a navigation-type task and there was significant delay duration variability, whereas in [16], the task was a navigation task in a simulated linear track and the delay duration variability was much smaller. Consistent with these results, we found more persistent responses in tasks with significant delay duration variability and in tasks with dynamic, motion-related inputs.

Our networks and learning paradigm had a number of biologically unrealistic features. Our networks consisted of simple generic rate neurons, whereas real neurons communicate via spikes and display a wide range of morphological and functional diversity. Moreover, our networks were trained with the biologically unrealistic backpropagation algorithm. However, a growing body of research demonstrates that task-trained generic neural networks like the ones we used in our simulations can capture many, sometimes surprisingly subtle, aspects of real biological circuits performing the same tasks [23, 37-39], implying that one may not always need highly biologically realistic architectures or learning rules to explain the behavior of complex neural circuits performing complex tasks. Our results contribute to this literature by showing that both sequential and nearly persistent, stable activity patterns experimentally observed in short-term memory studies are part of a spectrum that emerges naturally in generic neural networks trained on short-term memory tasks under different conditions.

## Methods

### Network details

We adopted a discrete-time formulation in which the network dynamics was described by:

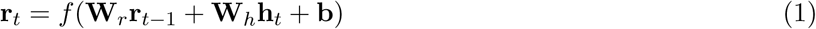

where **r**_*t*_ and **h**_*t*_ are the responses of the recurrent and input units at time *t*, respectively. Note that some previous studies start with a continuous-time formulation and obtain a discrete-time version through the Euler method. This yields an equation with the following form:

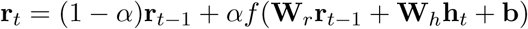

where *α* ≡ ∆*t*/*τ* describes the time step of the simulation in units of the intrinsic time scale of individual units. Typically, *α* is chosen to be small (e.g. *α* ~ 0.05 − 0.1), which is equivalent to assuming a long time constant for individual units. In contrast, our formulation (Equation 1) corresponds to taking *α* = 1, which does not make a long time constant assumption, but note that we increase the effective time constant of individual units through our initialization of **W**_*r*_ instead. More specifically, the hyperparameter *λ*_0_ controls the initial effective time constant of the units in our formulation. We set ∆*t* = 10 ms in all results reported in this paper.

For the main experiments, we used linear rectification (ReLU) for the nonlinearity *f*. All networks had 50 Poisson neurons in each input population and 500 recurrent neurons with ReLU activation. In networks with Hebbian synaptic plasticity, the general equation describing the network dynamics can be expressed as:

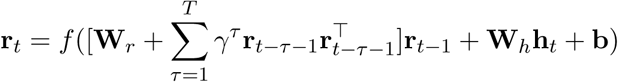

In practice, however, we found networks with *T* > 1 to be very unstable and difficult to train, hence we set *T* = 1, yielding the following equation:

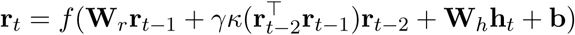

where *κ*(·) is a clipping function that clips its input between 0 and 100 to ensure stability and ϒ controls the strength of the Hebbian contribution. ϒ was set to 0.0005 in the change detection task and to 0.0007 in all other tasks. These values were the largest ϒ values that allowed the network to train successfully starting from at least 10 different initial conditions.

### Task details

In change detection, delayed estimation and gated delayed estimation tasks, we used circular stimulus spaces, which can be thought of as orientation, for example. The input neurons had von Mises tuning functions with circular means uniformly spaced between 0 and *π* and a constant concentration parameter *κ* = 2. The stimuli were drawn uniformly between 0 and *π*. In the 2AFC and comparison tasks, linear stimulus spaces were used. In the 2AFC task, the input neurons had Gaussian tuning functions with centers uniformly spaced between −40 and 40, and a constant standard deviation of 10. The stimuli presented were either −15 or 15 (randomly chosen in each trial) corresponding to the left and right choices, respectively. In the comparison task, the input neurons had Gaussian tuning functions with centers uniformly spaced between −50 and 50, and a constant standard deviation of 10. The stimuli were drawn uniformly between −40 and 40. In all tasks, during the stimulation periods, the gains of the input neurons were set to 1*/T*_stim_ at each time step (where *T*_stim_ denotes the duration of the stimulation period), yielding a cumulative gain of 1 for each input neuron throughout the stimulation period. All input neurons also had a stimulus-independent, uniform spontaneous gain of 0.1*/T*_delay_ at each time point during the delay, yielding a cumulative spontaneous firing rate of 0.1 spikes/s throughout the delay period. In all tasks, each trial took 1500 ms (150 simulation steps): 250 ms (25 simulation steps) for the stimulus period, 1000 ms (100 simulation steps) for the delay period and 250 ms (25 simulation steps) for the response period.

### Training details

The networks were trained with the Adam stochastic gradient descent algorithm [40] with learning rate 0.0005 and using the appropriate cost function for each task: mean squared error for continuous output tasks and cross-entropy for categorical tasks. For all tasks, we put an additional *l*_2_-norm regularizer (with coefficient 0.0001) on the mean activity of all recurrent units in the last 50 ms of each trial. In the tethering tasks, the coefficient of this regularizer was increased to 0.1. Batch size was 50 trials in all experiments. The networks were trained for 25000 iterations and tested on 300 new trials. All analyses were performed on these test trials.

### Analysis details

Ideal observer models for each task were derived based on earlier work (e.g. [36,44]) and the optimal performance was calculated from these ideal observers. As in [36], for the categorical tasks (COMP, CD, 2AFC), we measured performance in terms of the fractional information loss, which is defined as the average KL-divergence between the actual posterior and the network’s output normalized by the mutual information between the class labels and the neural responses. For the continuous output tasks (GDE, DE), performance was measured in terms of the fractional RMSE, which is defined as 100 × (RMSE_netw_ − RMSE_opt_)/RMSE_opt_, where RMSE_netw_ is the RMSE of the network and RMSE_opt_ is the RMSE of the ideal observer. In all the analyses presented in this paper, we only considered networks that had at most 50% information loss or fractional RMSE on the test set.

In calculating the sequentiality index (SI) for a given trial, we only included the recurrent neurons that had an average response of at least 0.1 during that trial. In addition, the entropy of the peak time distribution, which is one of the determinants of the SI, was calculated by dividing the total trial duration into 20 bins and calculating the Shannon entropy of the resulting count distribution. A pseudo-count of 0.1 was added to each bin before calculating the entropy.

In the simulated trials shown in Figure 6d-f, a randomly selected set of 100 recurrent units (out of 500 units) received unit inputs for the entire duration of the trial, while the remaining units did not receive any direct input.

## Data availability

The raw simulation data used for generating each figure are available upon request.

## Code availability

The code for reproducing the experiments and analyses reported in this paper is available at: https://github.com/eminorhan/recurrent-memory.

## Acknowledgments

This work was supported by Grant R01EY020958 from the National Eye Institute. We thank the staff at the High Performance Computing cluster at NYU, especially Shenglong Wang, for their help with troubleshooting. We thank an anonymous reviewer for suggesting the multi-task learning experiments.

---

**Figure S1:**
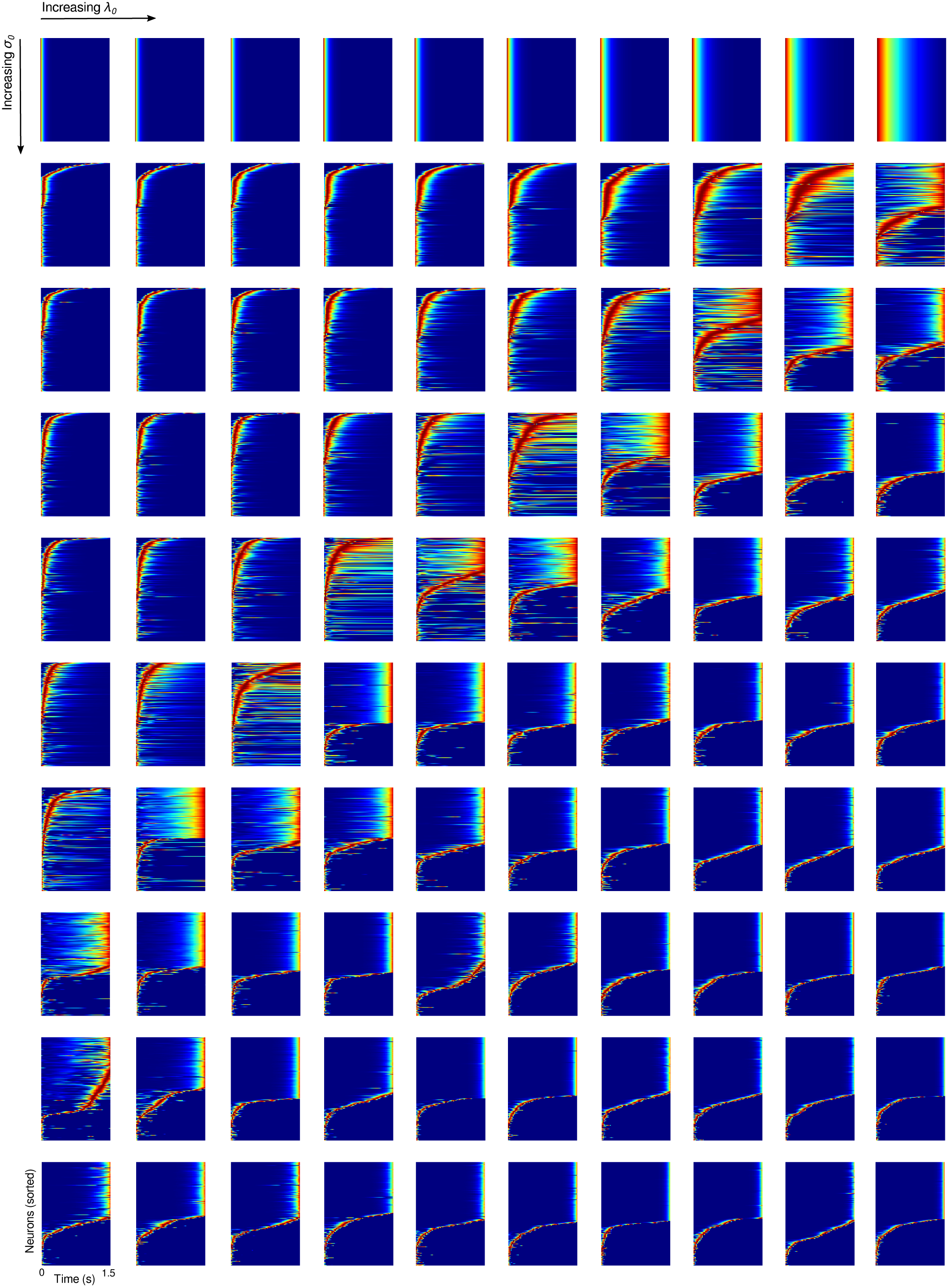
Initial, untrained network dynamics for different (*λ*_0_*, σ*_0_) values. The heat maps show the normalized responses of the recurrent units to a unit pulse delivered at time *t* = 0 to all units. Here, *λ*_0_ takes 10 uniformly-spaced values between 0.8 and 0.98 (columns) and *σ*_0_ takes 10 uniformly-spaced values between 0 and 0:4025 (rows).

**Figure S2:**
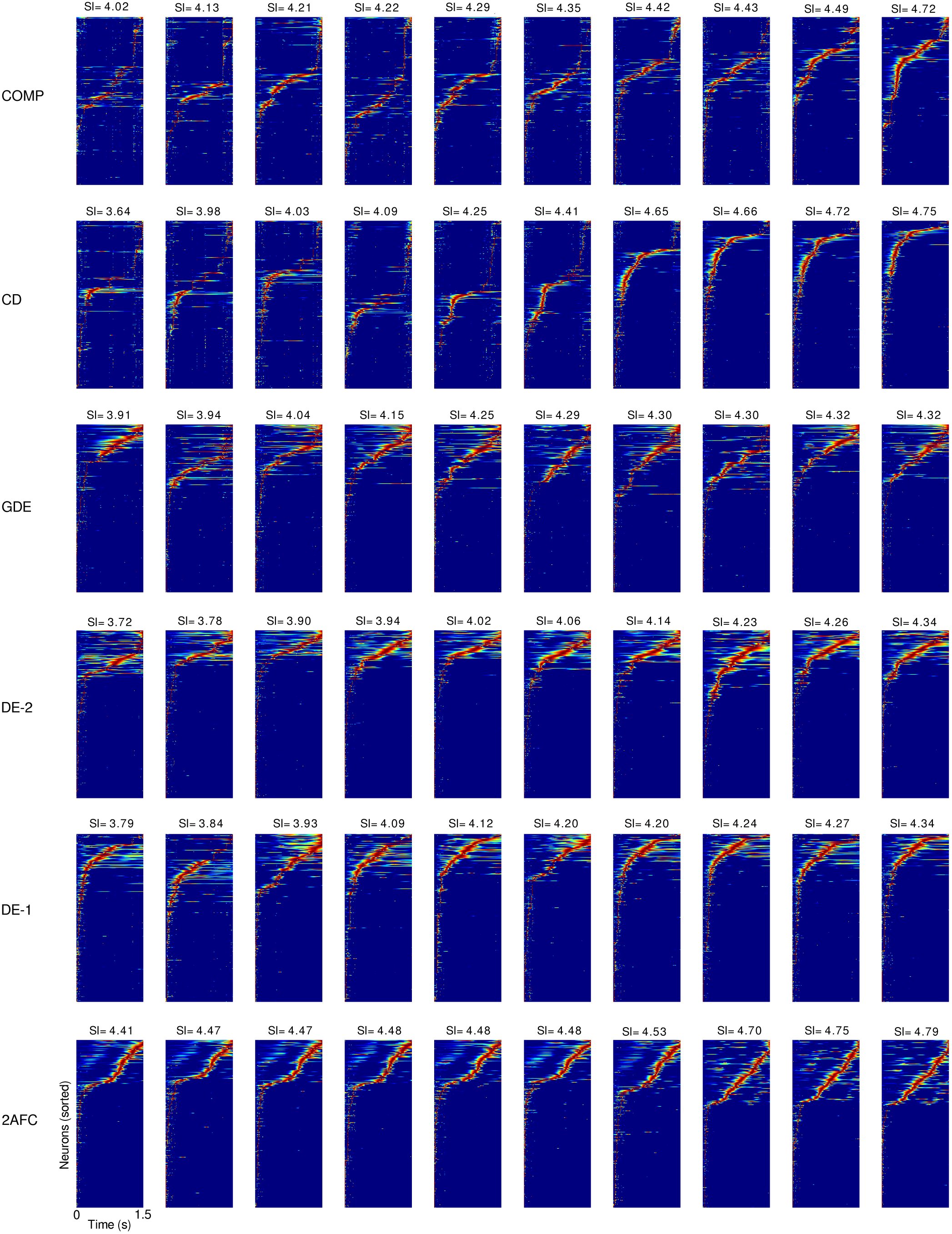
Example trials from the six tasks (basic condition). The SIs of the trials are indicated at the top of the plots. Trials are ordered by increasing SI from left to right. All trials shown here are from networks trained with *λ*_0_ = 0.96, *σ*_0_ = 0.313, *ρ* = 0. After training, all networks shown here achieved a test set performance within 25% of the optimal performance. In Supplementary Figures S2-S5, only the active recurrent units are shown.

**Figure S3:**
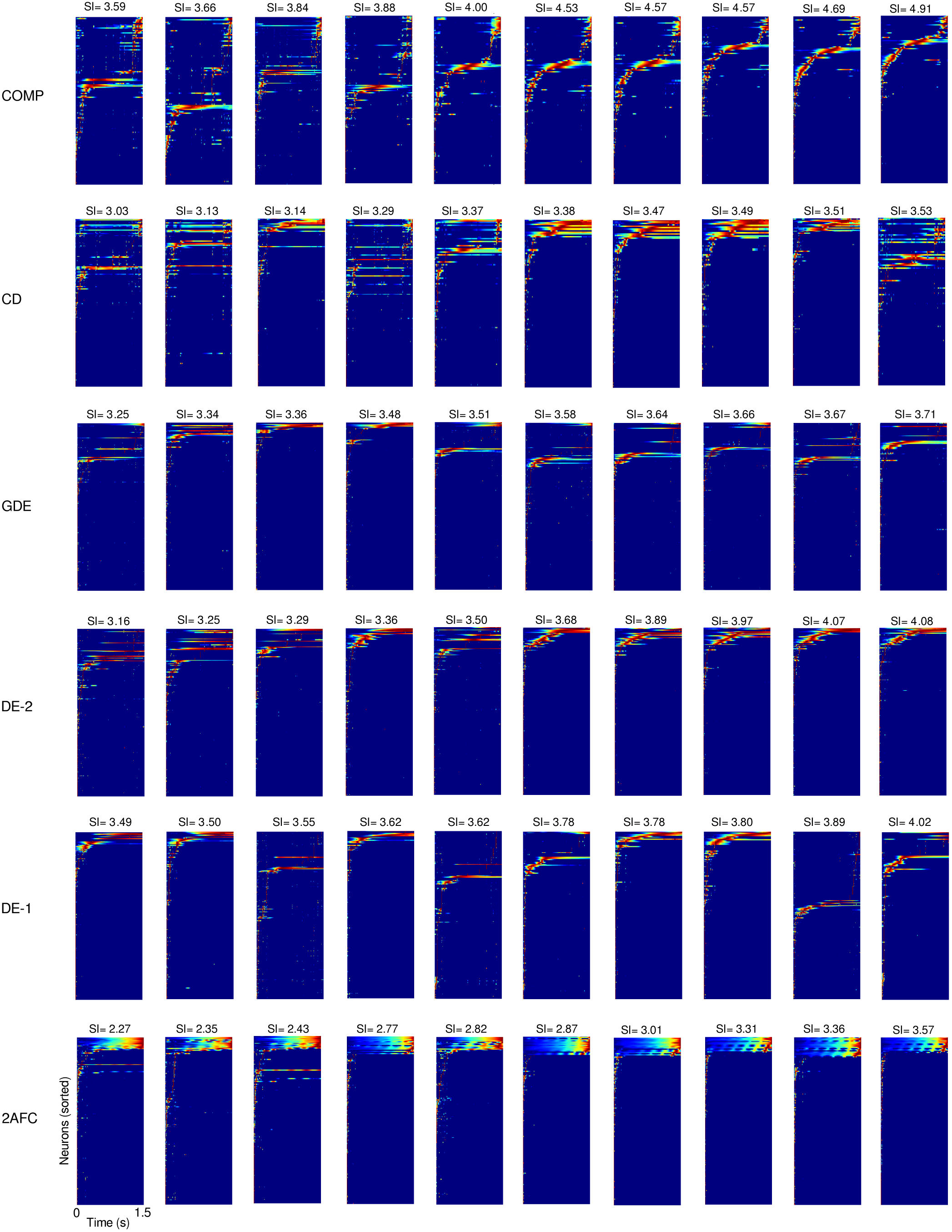
Example trials from the six tasks (basic condition). The SIs of the trials are indicated at the top of the plots. Trials are ordered by increasing SI from left to right. All trials shown here are from networks trained with *λ*_0_ = 0.96, *σ*_0_ = 0.134, *ρ* = 0. After training, all networks shown here achieved a test set performance within 50% of the optimal performance.

**Figure S4:**
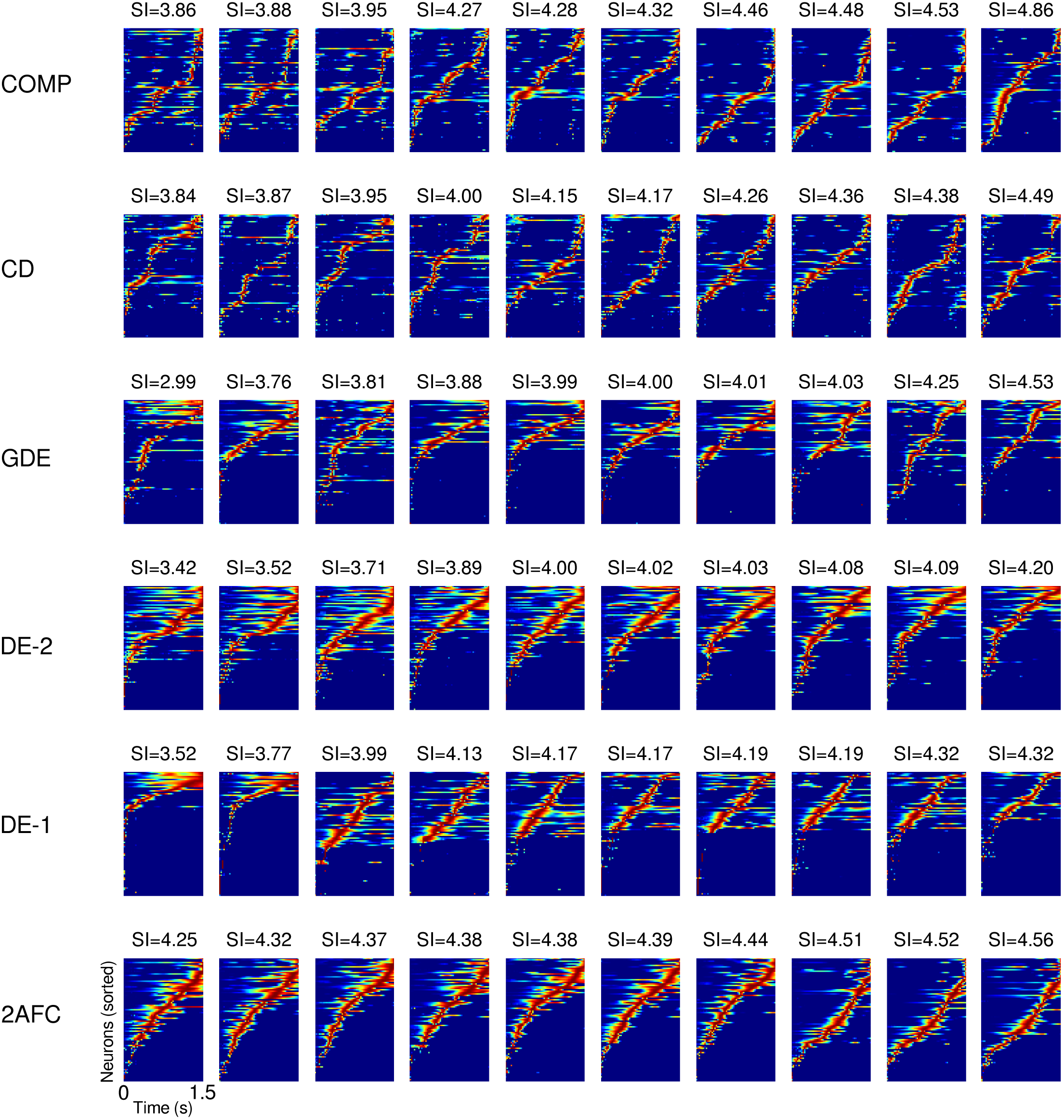
More example trials from the six tasks (basic condition). The SIs of the trials are indicated at the top of the plots. Trials are ordered by increasing SI from left to right. All trials shown here are from networks trained with *λ*_0_ = 0.96, *σ*_0_ = 0.313, *ρ* = 10^−3^. After training, all networks shown here achieved a test set performance within 50% of the optimal performance.

**Figure S5:**
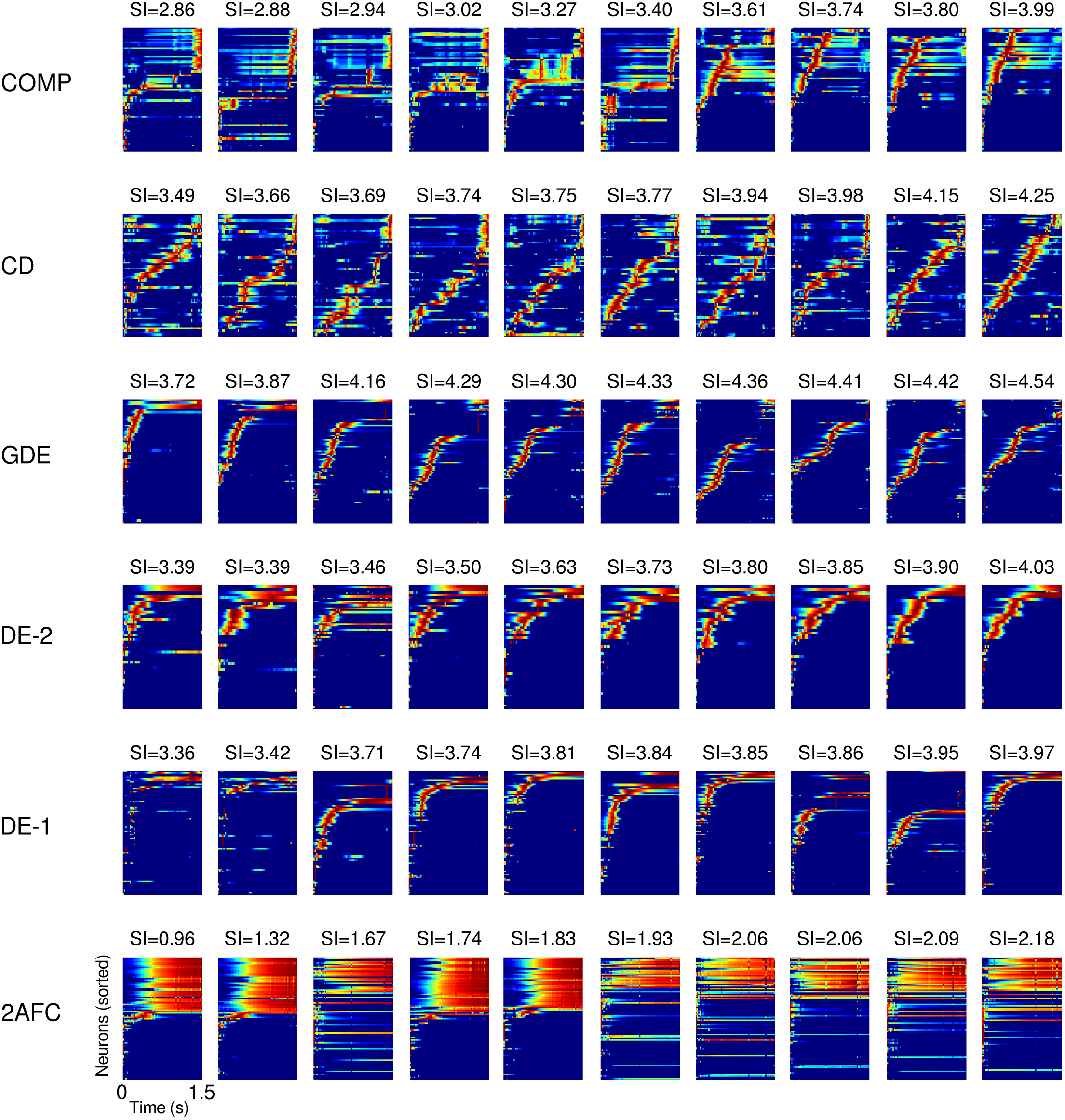
More example trials from the six tasks (basic condition). The SIs of the trials are indicated at the top of the plots. Trials are ordered by increasing SI from left to right. All trials shown here are from networks trained with *λ*_0_ = 0.96, *σ*_0_ = 0.134, *ρ* = 10^−3^. After training, all networks shown here achieved a test set performance within 50% of the optimal performance.

**Figure S6:**
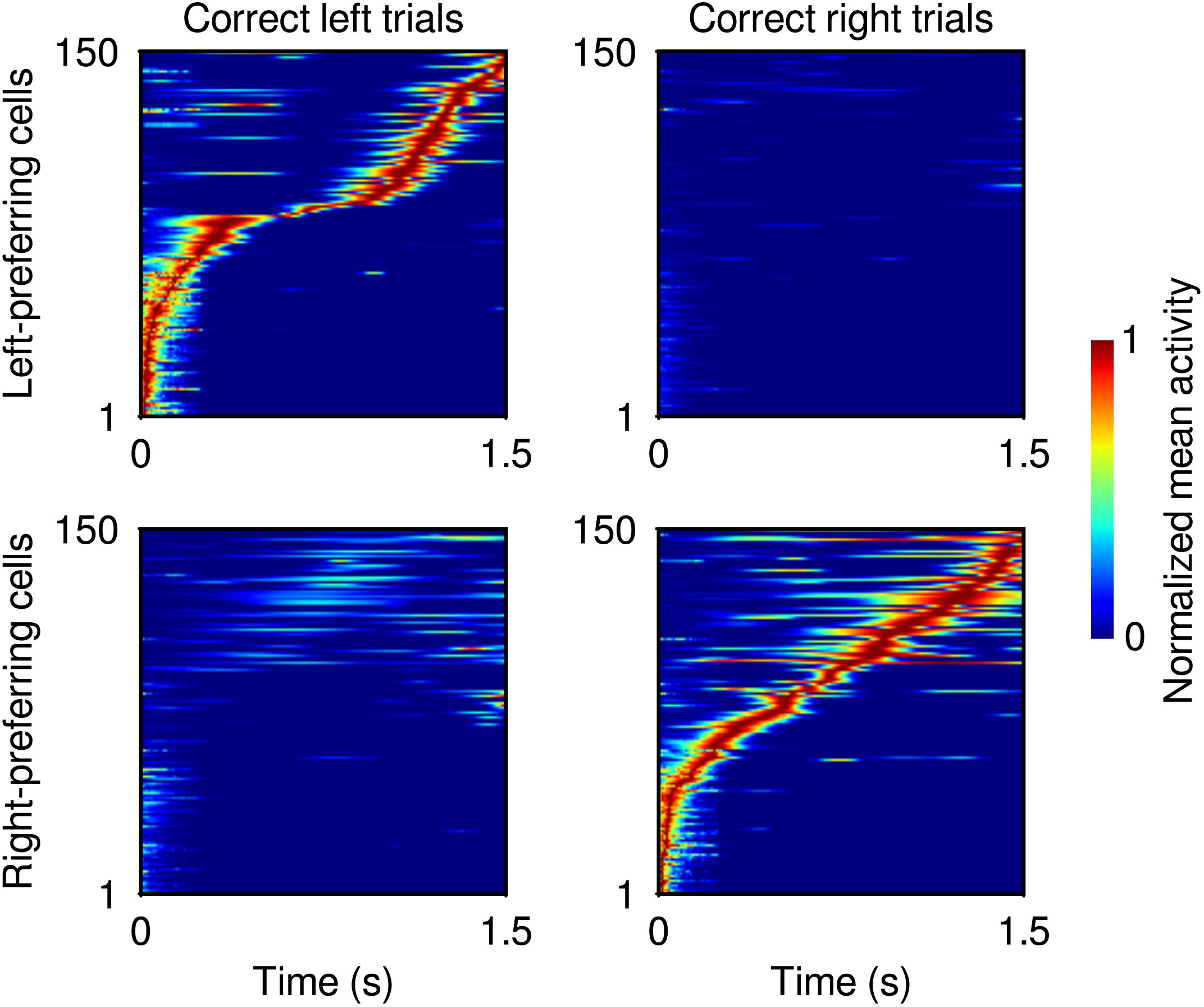
Average normalized activity of recurrent units in an example network trained in the 2AFC task. The network shown here was trained with *λ*_0_ = 0.96, *σ*_0_ = 0.313, *ρ* = 0. After training, the network achieved a test set performance within 0.1% of the optimal performance. As in ref. [16], we divided the recurrent units into left-preferring and right-preferring ones based on whether they responded more strongly during correct left choices or during correct right choices. The upper panel shows the average normalized responses of the left-preferring units in the correct left and correct right trials, respectively. Similarly, the lower panel shows the average normalized responses of the right-preferring units in the correct left and correct right trials. As reported in ref. [16], the trained network developed choice-specific sequences in the 2AFC task (cf. Figure 2c in ref. [16]). Only the most active 150 units from each group are shown in this figure; as always, the original network contained 500 recurrent units. This figure also demonstrates that the sequences are consistent from trial to trial, since the sequential activity pattern does not disappear when the responses are averaged over multiple trials.

**Figure S7:**
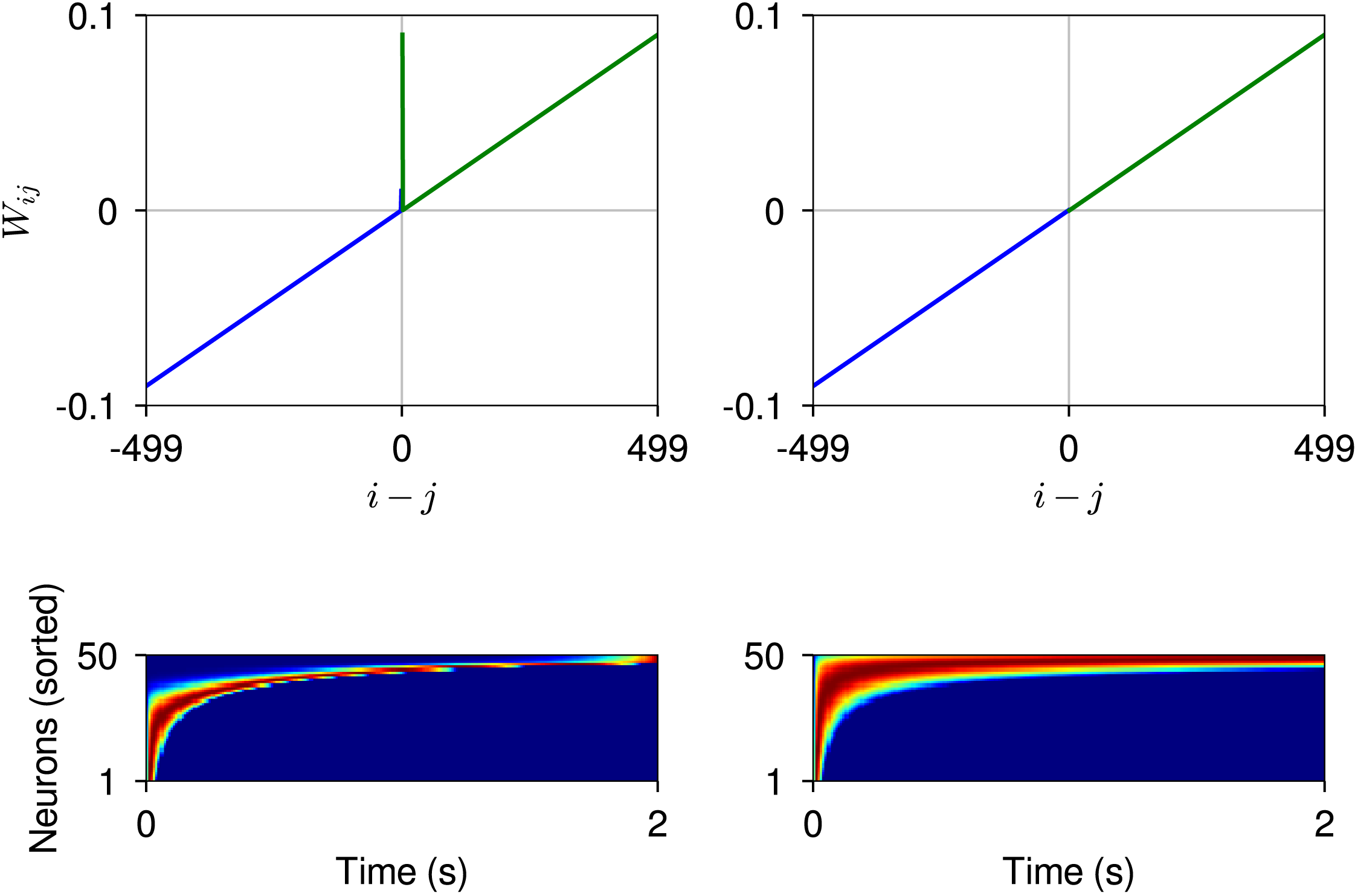
A simplified model that only incorporated the ReLU nonlinearity and the mean recurrent connection weight profiles shown in the upper panel (with no fluctuations around the mean) qualitatively captured the difference between the emergent sequential vs. persistent activity patterns (lower panel, left and right plots respectively). The networks simulated here had 500 recurrent units (only the most active 50 units are shown in the lower panel). All recurrent units received a unit pulse input at *t* = 0. The self-recurrence term in the recurrent connectivity matrix (not shown in the upper panel for clarity) was set to 1 in both cases. In the sequential case, the off-diagonal band was set to 0.09 in the forward direction and 0.01 in the backward direction, i.e. *W_i,i−_*_1_ = 0.09 and *W_i−_*_1,*i*_ = 0.01. The recurrent units did not have a bias term and they did not receive any direct inputs during the trial other than the unit pulse injected at the beginning of the trial.

**Figure S8:**
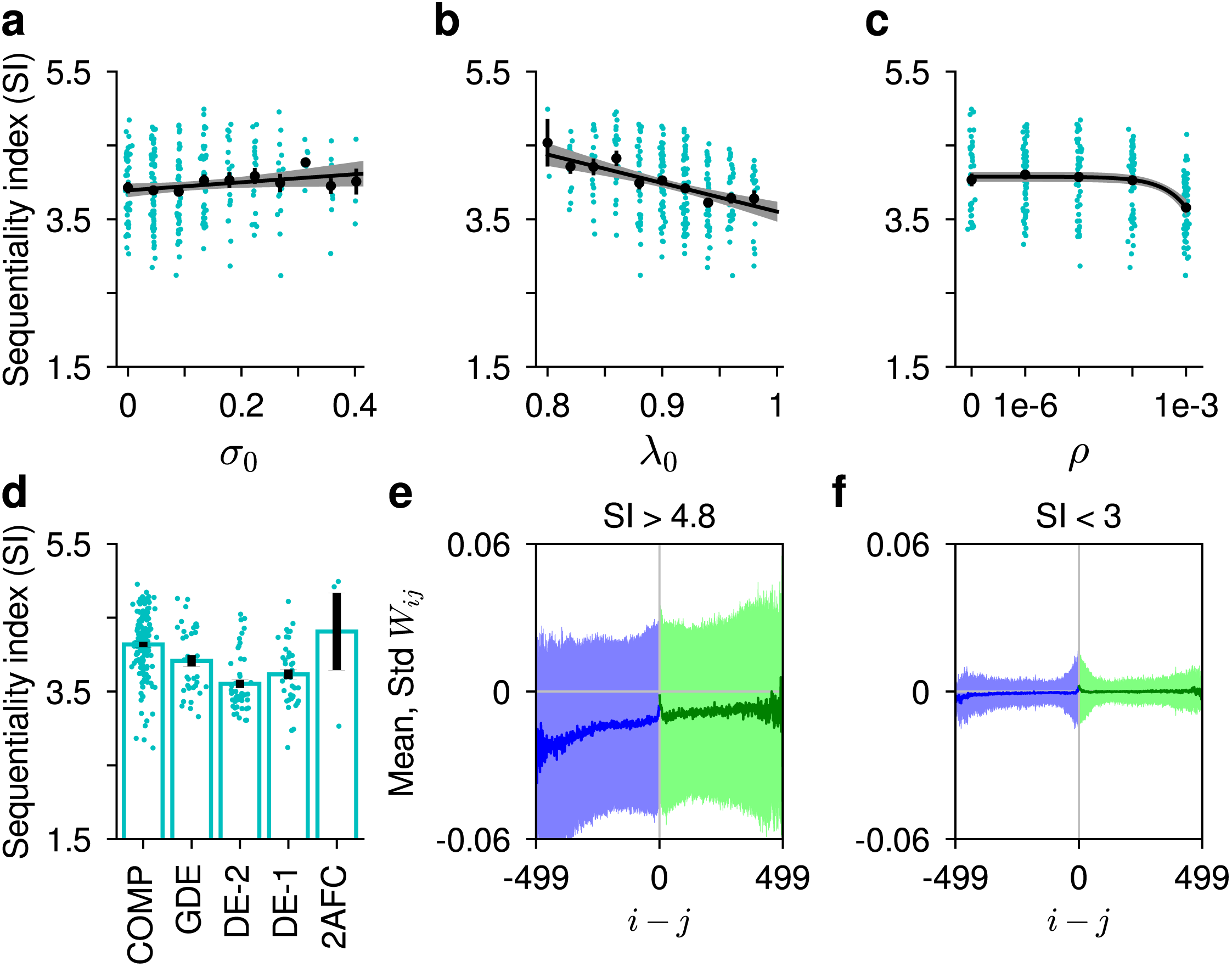
Results from the clipped ReLU networks. The clipped ReLU nonlinearity is similar to ReLU except that it is bounded above by a maximum value: i.e. *f* (*x*) = *clip*(*x, r_min_, r_max_*), where *r_min_* = 0 and *r_max_* = 100. **a** SI increased with *σ*_0_ (linear regression slope: 0.55, *R*^2^ = 0.01, *p* < .05). **b** SI decreased with *λ*_0_ (linear regression slope: −3.87, *R*^2^ = 0.11, *p* < 10^−7^). Note that this result differs from the corresponding result in the case of ReLU networks, where *λ*_0_ did not have a significant effect on the SI (Figure 2c). **c** SI decreased with *ρ* (linear regression slope: 418, *R*^2^ = 0.13, *p* < 10^−9^). **d** SI as a function of task. Overall, the ordering of the tasks by SI was similar to that obtained with the ReLU nonlinearity (Figure 3a). However, note that training was substantially more difficult with the clipped ReLU nonlinearity than with the ReLU nonlinearity. Across all tasks and all conditions, ReLU networks had a training success (defined as reaching within 50% of the optimal performance) of ~ 60%, whereas the clipped ReLU networks had a training success of only ~ 9.3%. In particular, we were not able to successfully train any networks in the CD task and very few in the 2AFC task. As a consequence, some of the differences between the tasks ended up not being significant in the clipped ReLU case. **e, f** Recurrent connection weight profiles (as in Figure 6a-c) in conditions where SI > 4.8 and in conditions where SI < 3, respectively. The weights were smaller in magnitude in **f**, because most of the low SI networks were trained under strong regularization.

**Figure S9:**
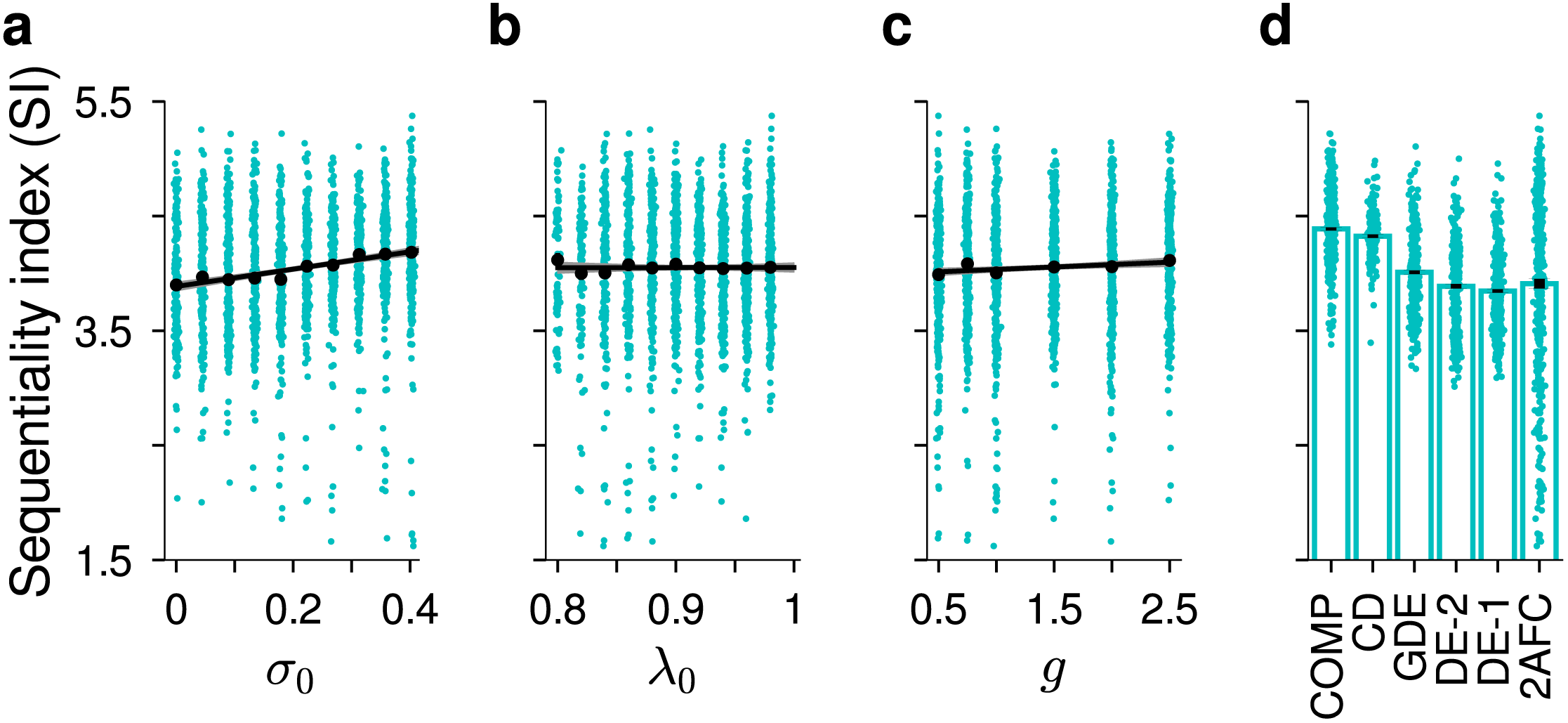
Changing the amount input noise. In these simulations, we set *ρ* = 0 and varied the gain of the input population(s), *g*. *g* = 1 corresponds to the original case reported in the main text; lower and higher values of *g* correspond to higher and lower amounts of input noise, respectively. **a** Combined across all noise conditions, SI increased with *σ*_0_ (linear regression slope: 0.76, *R*^2^ = 0.04, *P* < 10^−20^). **b** *λ*_0_ did not have a significant effect on SI (*p* = 0.96). **c** The input gain *g* slightly increased the SI (linear regression slope: 0.04, *R*^2^ = 0.003, *p* < 0.01). **d** Again, combined across all input noise levels, the ordering of the tasks by SI was similar to that obtained in the main set of experiments, where *g* = 1 (Figure 3a).

**Figure S10:**
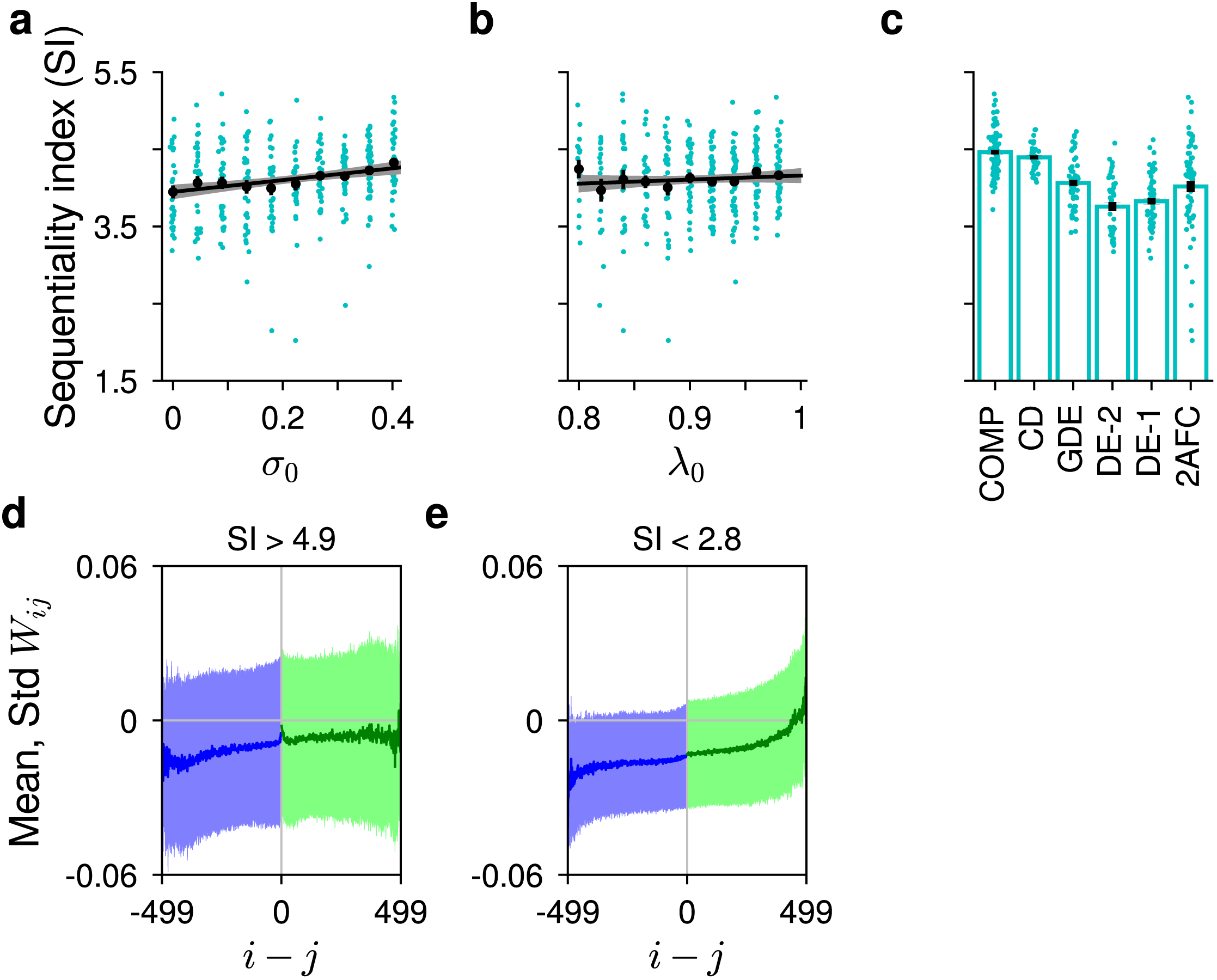
This figure shows the results when the analysis is restricted to the lowest level of input noise (*g* = 2.5). **a** SI increased significantly with *σ*_0_ (linear regression slope: 0.76, *R*^2^ = 0.05, *P* < 10^−4^). **b** *λ*_0_ did not have a significant effect on SI (*p* = 0.25). **c** The ordering of the tasks by SI was similar to that obtained in the main set of experiments. **d, e** Recurrent connection weight profiles (as in Figure 6a-c) in conditions where SI > 4.9 and in conditions where SI < 2.8, respectively.

**Figure S11:**
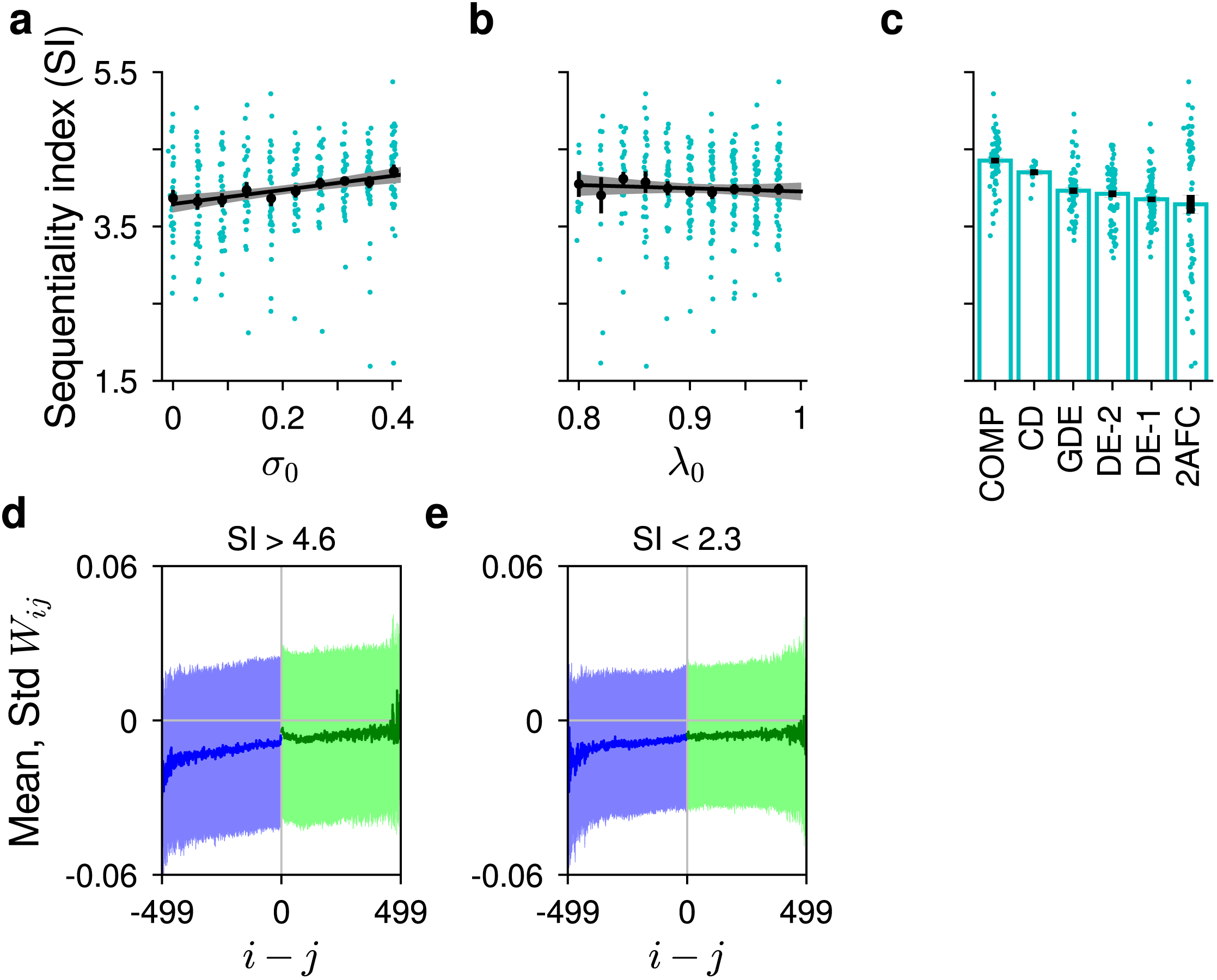
This figure shows the results when the analysis is restricted to the highest level of input noise (*g* = 0.5). **a** SI increased significantly with *σ*_0_ (linear regression slope: 0.91, *R*^2^ = 0.05, *P* < 10^−4^). **b** *λ*_0_ did not have a significant effect on SI (*p* = 0.46). **c** The ordering of the tasks by SI was similar to that obtained in the main set of experiments. **d, e** Recurrent connection weight profiles (as in Figure 6a-c) in conditions where SI > 4.6 and in conditions where SI < 2.3, respectively.

**Figure S12:**
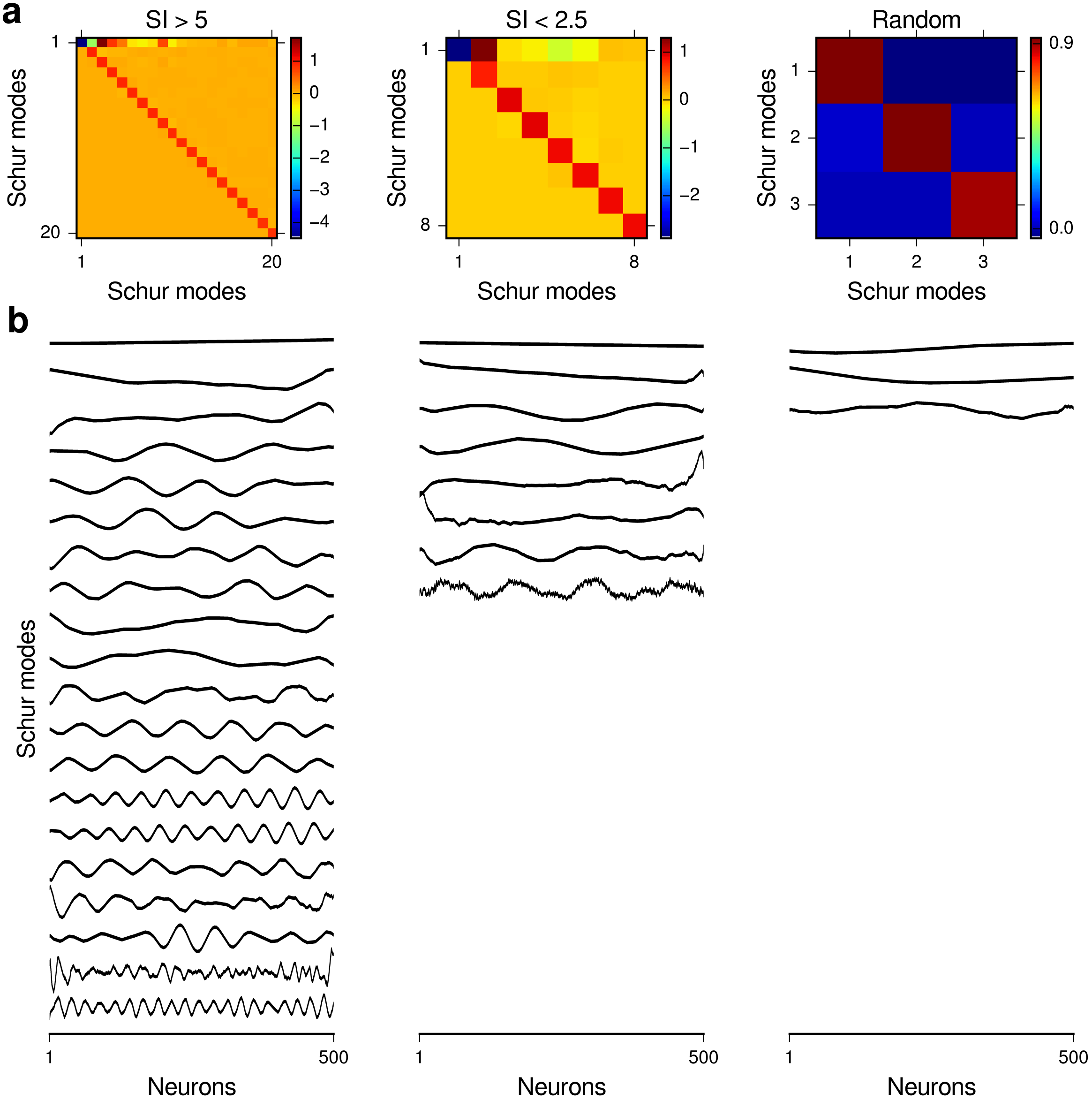
Schur decomposition of trained and random connectivity matrices. **a** Schur mode interaction matrices for the mean recurrent connectivity patterns shown in Figure 6a-c. Only significant Schur modes with at least one interaction of magnitude greater than 0.04 with another Schur mode are shown here. **b** The corresponding significant Schur modes. Networks with more sequential activity (SI > 5) have more high-frequency Schur modes than networks with less sequential activity (SI < 2.5). The random networks are close to normal.

**Figure S13:**
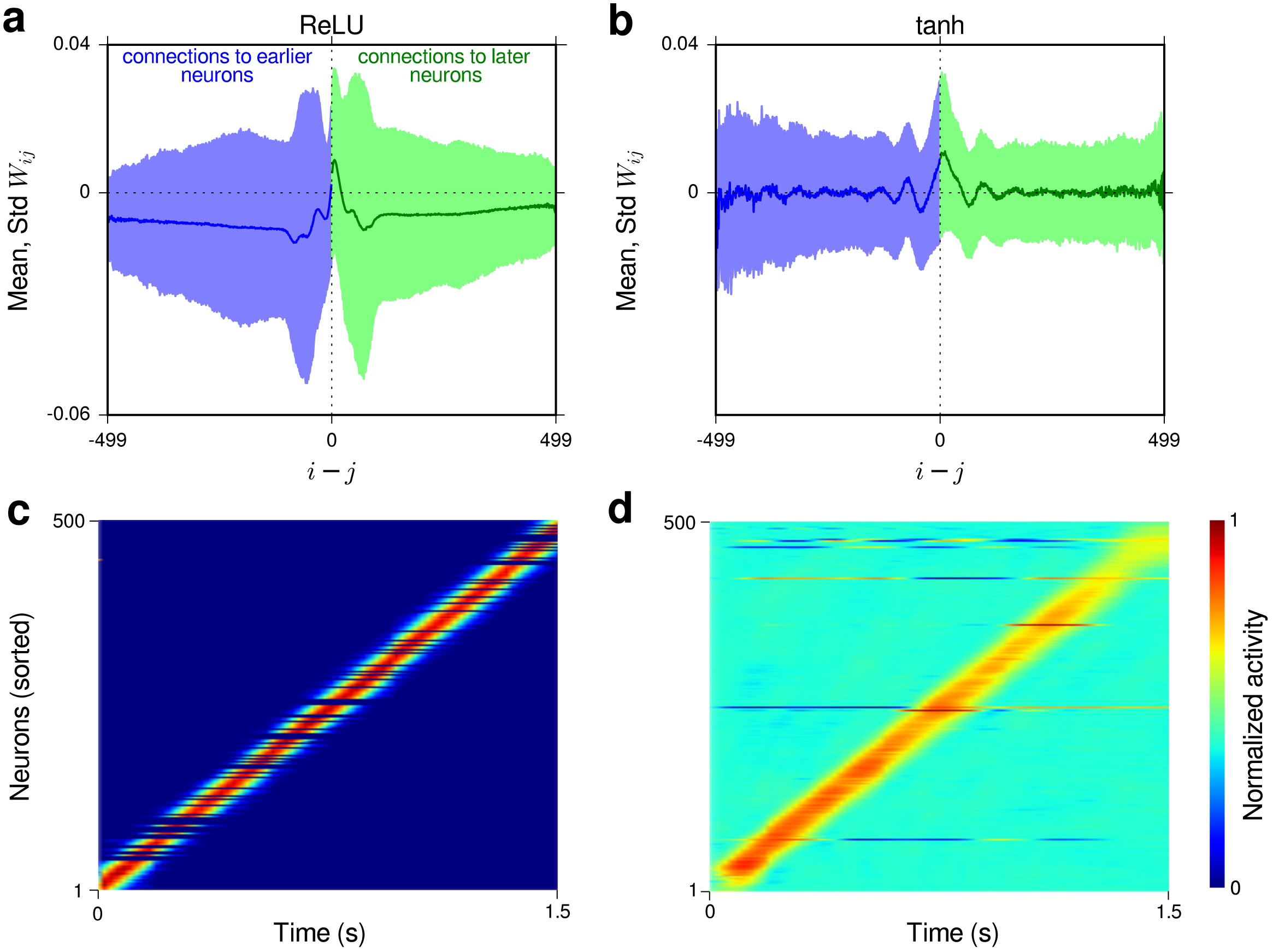
Results from networks explicitly trained to generate sequential activity as in ref. [35]. **a**-**b** are analogous to Figure 6a-b and show the recurrent weight profiles obtained in trained networks with ReLU and tanh nonlinearities, respectively. **c**-**d** show example trials for the corresponding networks (trained with the same initial condition). Only networks with sequentiality index larger than 5.45 were included in the results shown here.

**Figure S14:**
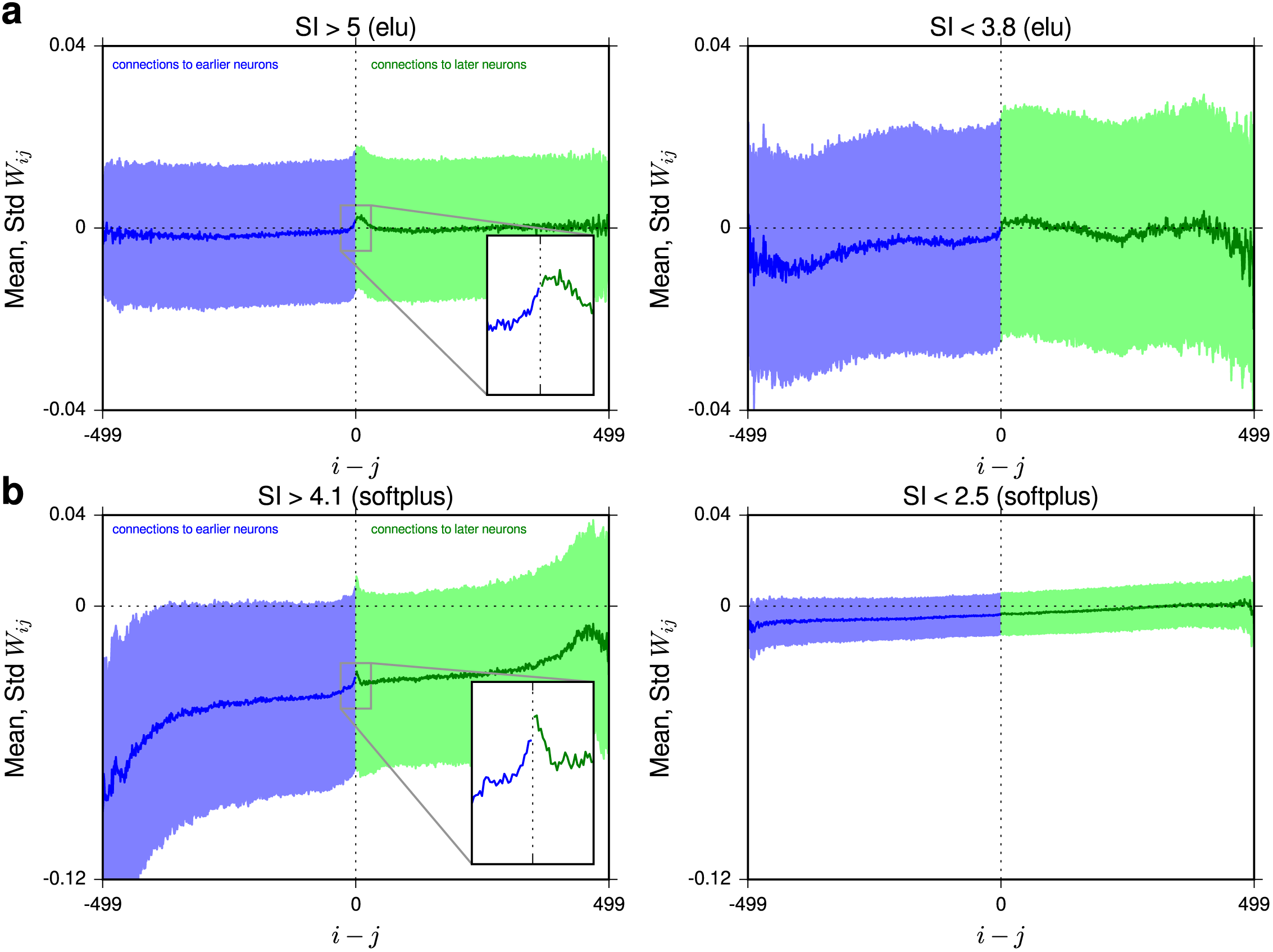
Circuit mechanism that generates sequential vs. persistent activity in networks with alternative activation functions. This figure is analogous to Figure 6a-b, but the results shown are for networks with the exponential linear (elu) activation function (**a**) and networks with the softplus activation function (**b**). Note that the elu activation function typically produced larger SIs than softplus, hence slightly different SI thresholds were used in the two cases to determine low and high SI networks.

